# Microbiome structure of a wild *Drosophila* community along tropical elevational gradients and comparison to laboratory lines

**DOI:** 10.1101/2021.07.28.454263

**Authors:** Joel J. Brown, Anna Jandová, Christopher T. Jeffs, Megan Higgie, Eva Nováková, Owen T. Lewis, Jan Hrček

## Abstract

Variation along environmental gradients in host-associated microbial communities is not well understood, compared to free-living microbial communities. Because elevational gradients may serve as natural proxies for climate change, understanding patterns along these gradients can inform our understanding of the threats hosts and their symbiotic microbes face in a warming world. In this study, we analysed bacterial microbiomes from pupae and adults of four *Drosophila* species native to Australian tropical rainforests. We sampled wild individuals at high and low elevation along two mountain gradients, to determine natural diversity patterns. Further, we sampled laboratory-reared individuals from isofemale lines established from the same localities, to see if any natural patterns are retained in the lab. In both environments, we controlled for diet to help elucidate other deterministic patterns of microbiome composition. We found small but significant differences in *Drosophila* bacterial community composition across elevation, with some notable taxonomic differences between different *Drosophila* species and sites. Further, we found that field-collected fly pupae had significantly richer microbiomes than laboratory-reared pupae. We also found similar microbiome composition in both types of provided diet, suggesting that the significant differences found among *Drosophila* microbiomes are the product of surrounding environments with different bacterial species pools, possibly bound to elevational differences in temperature. Our results suggest that comparative studies between lab and field specimens help reveal the true variability in microbiome communities that can exist within a single species.

**Importance:** Bacteria form microbial communities inside most higher-level organisms, but we know little about how the microbiome varies along environmental gradients and between natural host populations and laboratory colonies. To explore such effects on insect-associated microbiomes, we studied gut microbiome in four *Drosophila* species over two mountain gradients in tropical Australia. We also compared these data to individuals kept in the laboratory to understand how different settings changed microbiome communities. We found that field sampled individuals had significantly higher microbiome diversity than those from the lab. In wild *Drosophila* populations, elevation explains a small but significant amount of the variation in their microbial communities. Our study highlights the importance of environmental bacterial sources for *Drosophila* microbiome composition across elevational gradients, and shows how comparative studies help reveal the true flexibility in microbiome communities that can exist within a species.

## Introduction

Patterns of diversity over environmental gradients like latitude, elevation, or environmental degradation have long been of interest in community ecology and are of renewed interest for studying the potential consequences of climate change^1–5^. Most studies have focused on animals and plants to investigate these patterns, but bacterial communities are receiving increased attention. Some studies suggest free-living bacteria do not follow the same broad biogeographic patterns as plants and animals^6–8^. Fierer et al.^1^ showed that soil bacteria did not change significantly in diversity when sampled across an elevational gradient, in contrast to trends documented in most other taxa. Subsequent studies have found inconsistent patterns in bacterial communities sampled from streams and soils across elevational gradients, with differences usually being attributed to changes in pH and C:N ratio ^2,8–10^.

Many insects maintain intimate communities of symbiotic microbes (their ‘microbiome’). Insect microbiomes can play important roles in host health, digestion, thermal regulation, and protection against natural enemies (reviewed in^11–13^). In turn, many factors can influence insect microbiome composition, some host-dependent (e.g. diet, insect species identity, ontogeny, and parent-to-offspring transmission), others host-independent (e.g. abiotic factors like local environment and temperature)^14–22^. Symbioses between insects and bacteria have been particularly well investigated^23^, notably because insect microbiome communities tend to be less complex than those of vertebrates^24^. However, in contrast to environmental microbial communities, the effect of elevational change on insect-associated microbiomes has yet to be investigated in-depth. The most conspicuous aspect of a change in elevation is a difference in mean temperature, creating different environments that can be used as a proxy for climate change scenarios^25,26^. Elevational differences in temperature mean we would expect to see differences in microbiome composition related to temperature-dependent development in both insects^27–31^ and bacteria^32–34^. Thus, at different elevations, and in climate change scenarios, insect-associated microbiomes could have different composition^20^ with potentially important consequences on microbiome and host functionality.

*Drosophila* spp. are established models for studying insect-associated microbiomes^35–40^ because they are cosmopolitan, occurring in a wide variety of habitats, and easy to maintain in laboratory cultures. *Drosophila*-associated microbiomes have important functional impacts on many aspects of their ecology including thermal tolerance^41^, development^42^, ability to recognise kin^43^, and immunity^38,44^. The microbiomes are of moderate-to-low diversity, making them relatively simple to characterise. Additionally, some *Drosophila* species/populations possess intracellular bacterial symbionts (*Wolbachia* and *Spiroplasma*) that can influence host immunity and protect against natural enemies, including pathogenic fungi, nematodes, and parasitoids^45–49^. This, combined with the well-studied nature of *Drosophila*, makes them ideal candidates for investigating insect-associated microbiomes over elevational gradients.

Here we examine the effects of elevation change on insect microbiome composition by focusing on the underlying biotic and abiotic factors, including elevation, host species identity, and site location. We sampled wild populations of four focal species of frugivorous *Drosophila* from two mountain gradients in Queensland, Australia: *Drosophila rubida, D. pseudoananassae, D. pallidifrons*, and *D. sulfurigaster*. These species occur throughout north Queensland along multiple altitudinal gradients in the Wet Tropics. We chose these four species because they are abundant and occur in sympatry across the full elevational gradient at our focal sites^50^. We hypothesised that variation in microbiome composition between high and low elevation populations will reflect temperature differences at these sites. To control for diet in the field we exclusively sampled pupae from banana-baited bottle traps (see Jeffs et al.^50^). The sampling approach guarantees that each analysed individual originated from an egg laid in our bottle traps and therefore fed solely on yeasted banana. To reinforce our investigation, we analysed lab-reared flies of the same species collected from the same field sites to test if their microbiomes retained any natural differences when reared in the laboratory on a standard, yeast-based diet. We expected *a priori* to find high among-individual variation and hypothesised that species identity, elevation, and environment of origin (i.e. lab vs field) would be the primary causes of difference in host microbiome composition^35,39,40,51^.

## 2. Material & Methods

### 2.1 Study Sites

The Australian Wet Tropics World Heritage Area is a 450 km long, narrow section of rainforest along Queensland’s northeast coast between Cooktown and Townsville (15-19’S, 145-146.30’E). Samples were collected from two altitudinal gradients: Paluma Range Road (within Paluma Range National Park 19°00’S, 146°14’E) and Kirrama Range Road (within Girramay National Park 18°12’S, 145°50’E). The Paluma gradient ranges from 59 m to 916 m above sea level (a.s.l.) and the Kirrama gradient ranges from 92 m to 770 m a.s.l^50^. We chose sites at high and low elevations (Paluma: 880m, 70m; Kirrama: 730m, 70m) to capture a ∼5°C temperature range (mean temperatures 21°C at high elevation, 26°C at low elevation^50^. Temperature was recorded by multiple dataloggers suspended next to bottle traps at each site, which took a reading every hour. Our previous study on these gradients determined ∼5°C temperature range and found substantial differences in *Drosophila* community composition at high and low elevation, with some species not present at the low and high ends of the gradient and others changing in abundance^50^.

### 2.2 Sample Collection and Selection

To ensure a comparable dataset, we selected stratified subsets of samples from the field and laboratory. We primarily collected pupae because they are more easily standardized than larvae and allow us to account for discrepancies in development rate between species. We additionally collected a small number of adult flies to compare microbiome composition between pupae and adults. Based on the results of Jeffs et al.^50^, which used pupae samples to identify the natural *Drosophila*-parasitoid food web with COI metabarcoding and Multiplex PCR methods, we selected 214 field samples of the four most abundant *Drosophila* species that occurred at all elevations along both transects: *Drosophila rubida, D. pseudoananassae, D. pallidifrons*, and *D. sulfurigaster* (Table 1). Eight *D. rubida* pupae were parasitised by parasitoid wasps, enabling us to test if there are any changes in microbiome richness or unique microbial taxa associated with a developing parasitoid. We subsequently sampled 70 pupae and 70 adults from isofemale (IF) laboratory lines (2-4 lines per species) of these four elevationally ubiquitous species (20 pupae and 20 adults from *D. sulfurigaster, D. rubida*, and *D. pseudoananassae*, and 10 pupae and 10 adults from *D. pallidifrons*) to investigate if suspected natural patterns (site- and species-specific influence) were retained in lab-reared flies. Additionally, we took 10 samples of the food source provided to lab-reared *Drosophila* and 20 samples of the banana bait we used in our field sampling, to compare *Drosophila* microbiome samples to a dietary reference and determine how congruent the microbiome communities were between the food source and the insect hosts. Table 1 presents a detailed breakdown of all samples used in this study.

**Table 1:**
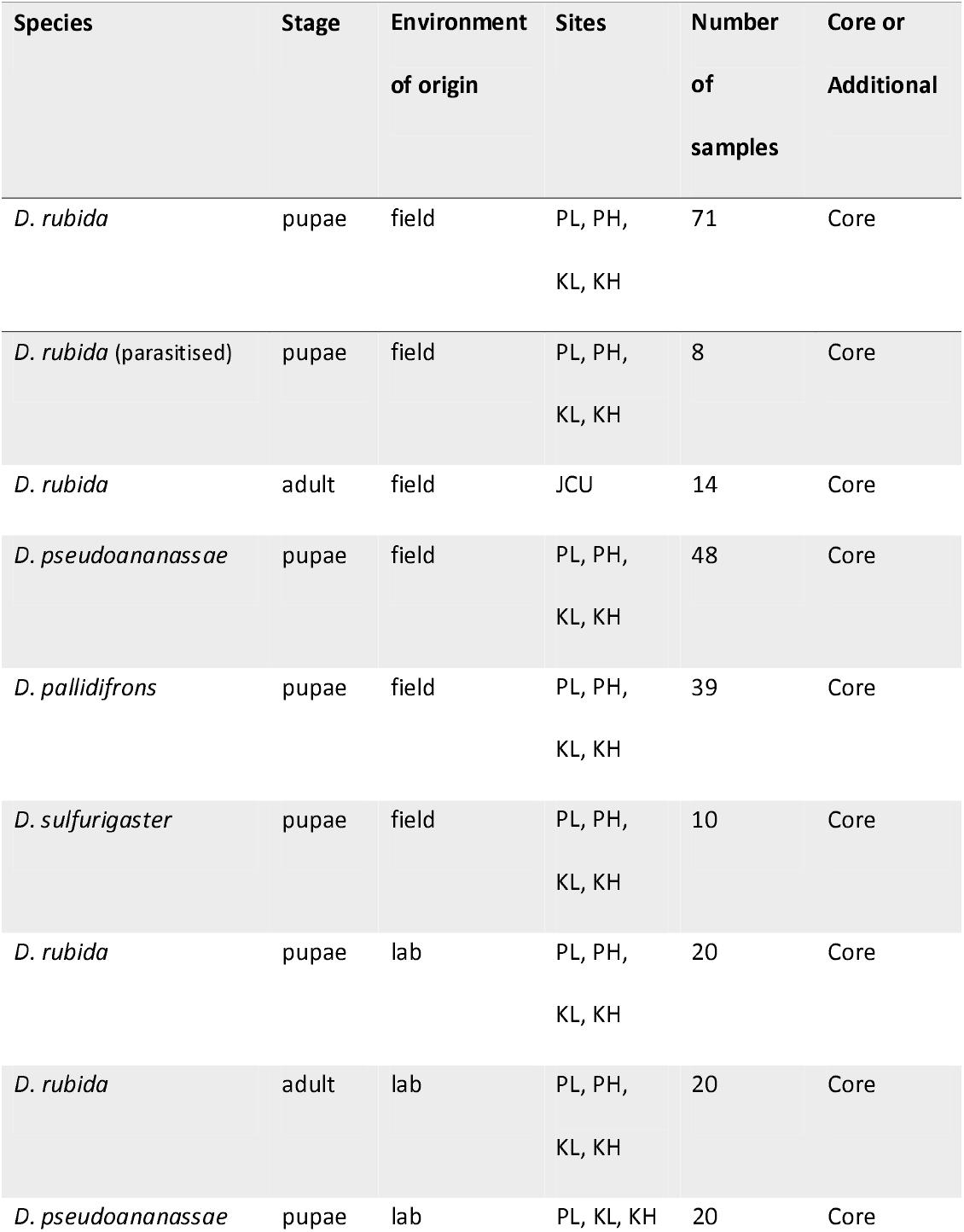
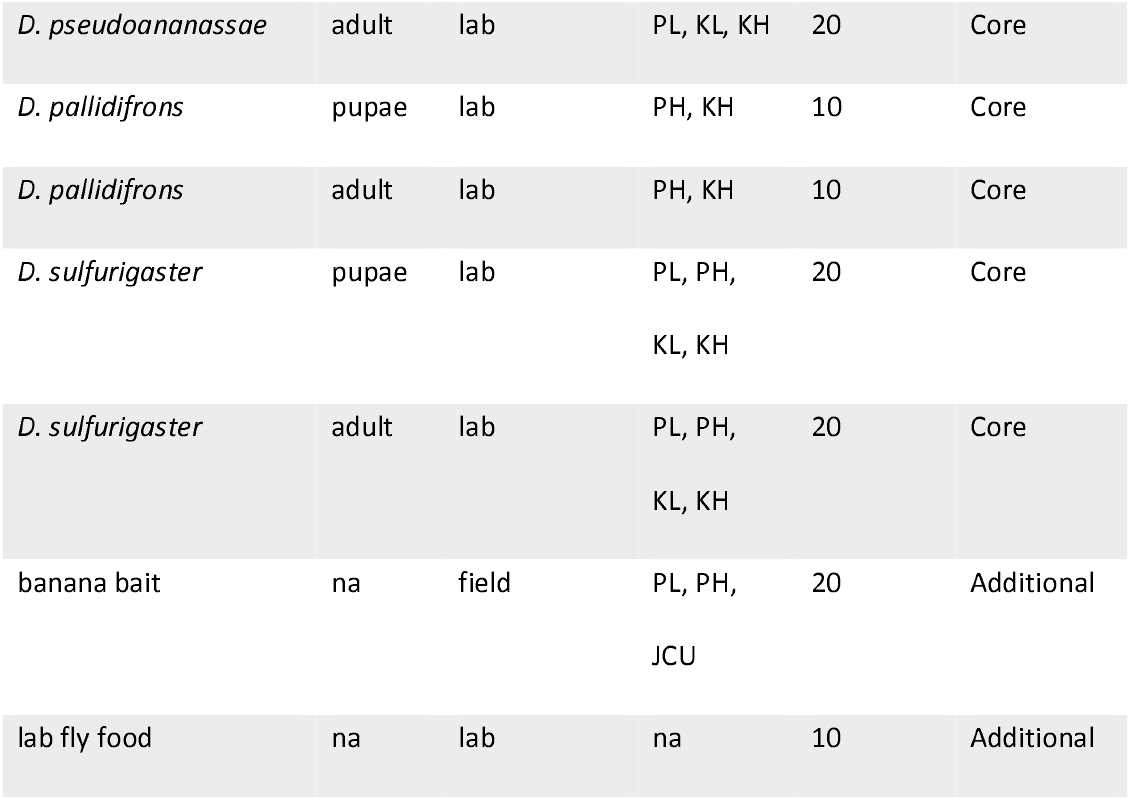
Breakdown of the sample set used in this study. PL = Paluma Low, PH = Paluma High, KL = Kirrama Low, KH = Kirrama High, JCU = James Cook University campus in Townsville.

The *Drosophila* pupae field samples were collected from banana-baited bottle traps placed at low and high elevation sites along both altitudinal gradients. Each bottle trap had a piece of cardboard to assist *Drosophila* larvae in pupation. Bottle traps were left exposed for either 11-12, 14-15, or 24 days, to capture the natural variation in community colonisation and ontogenetic development in different *Drosophila* species^50^. On the day of sampling, these cards were removed and sealed in ziplock bags. The individuals we collected as pupae thus only fed on banana bait. Pupae from each card were sampled by placing the card on a white plastic dinner plate and adding distilled water, using a small paintbrush to remove all pupae. Each pupa was placed into an individual well in 96-well PCR plates and preserved in 100% ethanol. Adults were aspirated from bottle traps hung at James Cook University, Townsville (JCU), 2 days after provision of fresh banana bait and placed into individual vials in 100% ethanol. JCU became a supplementary sampling site after Kirrama became inaccessible due to heavy rainfall and landslides. Laboratory-reared pupae were collected with forceps from standard fly food. Adults were collected with an aspirator and sexed, then placed in individual vials with 100% ethanol. Laboratory isofemale lines were established from the same populations sampled in the field (i.e., they were collected at the same sites and shipped live to the lab in Czech Republic in 2017-18, after collection of field samples in 2016). Isofemale lines were kept in the lab on a standard *Drosophila* diet medium (corn flour, sugar, agar, yeast, and methyl-4-hydroxybenzoate) for between 18-30 months by the time of sampling.

Sample DNA was individually extracted using single column GeneAid Blood and Tissue kits according to manufacturer instructions, with one extraction negative control accompanying every 29 samples. For confirmation of *Drosophila* species identification we utilised a multiplex PCR approach^50^. The PCRs were based on the ITS2 and COI regions using custom specific forward and reverse primers. We combined primers into multiplex PCR reactions based on difference in length of product. Gel electrophoresis of PCR products showed product length, allowing us to identify species. When multiplex PCR was inconclusive, we used Sanger sequencing with custom Diptera specific primers to identify the *Drosophila* species. Full details of these processes and primer sequences can be found in the Supporting Information of ^50^.

### 2.3 Library Preparation & Sequencing

After extraction and identification, all samples were moved to 96-well plates in a randomised order in preparation for bacterial sequencing. DNA templates were stored at - 75ºC. These templates were used for amplification of ∼400 bp of the V4/V5 hypervariable region of the 16S rRNA gene according to Earth Microbiome Project standards (EMP; http://www.earthmicrobiome.org/protocols-and-standards/16s/). Briefly, sample multiplexing was based on the EMP-proposed double barcoding strategy using the EMP-recommended modifications (12 bp Golay barcodes included on the forward primer 515F (5⍰-GTGYCAGCMGCCGCGGTAA), and additional 5 bp barcodes on the reverse primer 926R (5⍰-CCGYCAATTYMTTTRAGTTT)^52,53^. We also added a custom 18S rRNA gene blocking primer (named 926X - 5⍰ GTGCCCTTCCGTCAATTCCT-C3 3’) to counteract the low specificity of EMP primers towards the 16S rRNA gene^22^. PCRs were carried out in triplicate and successful amplification was confirmed with gel electrophoresis. Combined triplicates for each sample were purified with AMPure XP (Beckman Coulter) magnetic beads and equimolarly pooled to a single library (based on DNA concentration measured using a Synergy H1 (BioTek) spectrophotometer). The library was purified using Pippin Prep (Sage Science) from all fragments outside the 300-1100 bp range. To control for contamination, PCR biases and confirm barcoding success, we included four negative controls from the extraction procedure (ENC), eight negative controls from the PCR process (NC), and eight positive controls (PC) of mock microbial communities. PCs were supplied commercially and comprised 4 samples of gDNA templates with equal abundance of 10 bacterial species (ATCC® MSA-1000™) and 4 samples with staggered abundance for the same bacteria (ATCC^®^ MSA-1001™). Altogether the library comprised PCR products from four 96-well plates, each containing one ENC, two NCs, and two different PCs. The library was sequenced in a single run of Illumina MiSeq platform using v3 chemistry with 2 × 300 bp output (Norwegian High Throughput Sequencing Centre, Department of Medical Genetics, Oslo University Hospital).

### 2.4 Data Processing

The sequencing process returned 15,893,914 reads. These raw reads were quality checked (FastQC^54^) and trimmed using USEARCH v9.2.64^55^, to keep the quality score above Q20. We trimmed the primers, demultiplexed and merged the reads which resulted in a final amplicon length of 357 bp. We then clustered the reads at 100% identity for a representative set of sequences and used the USEARCH global alignment option at both 99% and 97% identity^55^ for *de novo* OTU assignment. We subsequently used the BLAST algorithm^56^ on the representative sequences, matching them against the SILVA 132 database^57^ for taxonomic identification, producing a dataset of 1132 OTUs at 97% identity and 1118 at 99% identity. We used the 97% identity OTU table as the primary dataset and the 99% identity table as a supplemental dataset to confirm that the patterns we found were not a product of identity threshold (Fig. S1). Raw sequence data is available on NCBI Raw Sequence Archive (accession no.: PRJNA849960).

We recovered a mean of 16,898 reads per sample and a median of 14,751 reads (not including negative controls). From our positive controls, we recovered microbiome profiles that matched the expected community composition in each of the ‘staggered’ and ‘even’ mock communities. The two low abundant species from the staggered templates (present at 0.04%) were successfully recovered from all four staggered mock samples. In the even mocks, there was consistent overrepresentation of *Clostridium beijerinckii* and *Escherichia coli* (1.4x - 4.7x expected), leading to reductions in other taxa. Overall, the positive controls in this sequencing run matched our previous sequence outcomes^22,58^.

Any chloroplast, mitochondrial, or eukaryotic OTUs were identified in the OTU table and excluded. Potential bacterial contaminants were systematically evaluated using the R package ‘decontam’ (V1.5.0^59^). The ‘decontam’ package uses the prevalence or frequency of OTUs detected in negative controls to remove suspected contaminants. We used the prevalence method, which compares the presence/absence of each sequence in true positive samples compared to the prevalence in negative controls. We used a strict threshold of 0.5, meaning any OTU with a higher proportion of reads in negative controls than test samples was excluded as a contaminant. The prevalence of each OTU is shown in Fig. S2. 43 OTUs were eliminated from the dataset via this process. Singletons were also excluded, as was any OTU that made up less than one percent of the total reads for each sample, which collectively removed 837 OTUs. These procedures resulted in a dataset of 110 OTUs and 360 samples.

### 2.5 Statistical Analyses

Sample analysis was carried out using the packages ‘vegan’^60^ and ‘phyloseq’^61^ in R^62^. To measure alpha diversity we calculated the Shannon diversity index, using paired Wilcoxon and paired ANOVA tests to compare values between sets of samples. We calculated Bray-Curtis dissimilarity as a quantitative measure of beta diversity and used these values to create non-metric multidimensional scaling ordinations (NMDS), to simultaneously evaluate the roles of elevation, site, *Drosophila* species, life stage, environment of origin, parasitism, trap identity, duration of trap exposure (and isofemale line and generation number for laboratory samples). To support these ordinations statistically, we calculated PERMANOVA tests on Bray-Curtis dissimilarity values. We applied the Benjamini-Hochberg (B-H) correction on PERMANOVA p-values to control for multiple comparisons and false discovery rate. We also evaluated statistical significance of differences in dispersion among groups using multivariate homogeneity analysis (function “betadisper” in ‘vegan’^60^) and ran each of these tests with 999 permutations. In addition, we used DeSEQ differential abundance analysis from the R package ‘microbiomeSeq’^63^ to compare the relative abundances of bacterial taxa across different sample sets, e.g., differential abundance between *Drosophila* species or between sites.

To systematically order our analyses, we first tested every variable (environment of origin, elevation, gradient identity, species identity, developmental stage, trap identity, trap duration of exposure, generation number, isofemale line) with all samples to determine the relative importance of all studied factors (results shown below). We subsequently separated the data by environment of origin, i.e., we tested lab *Drosophila* samples separately and field *Drosophila* samples separately. Splitting the dataset in this way meant some variables were not relevant to field samples, and vice versa. For example, when analysing field-only samples we did not include isofemale line or generation number because these only pertained to lab samples. Similarly, when focusing on lab-only samples, we did not include trap identity, trap exposure time, or parasitoid status, because none were parasitised, and traps were only a component of the field study. Furthermore, trap identity and site were correlated variables because each trap was only used at one site. We tested the food/bait samples separately because they were not focal samples and were used to support conclusions about *Drosophila* microbiomes. The only samples positive for parasitoid detection were from *Drosophila rubida* pupae, thus we only compared these samples to other *D. rubida* pupae, and not as part of the core analyses.

## 3. Results

We first tested which of the studied factors influenced microbiome composition in all core field and lab samples combined, using PERMANOVA analysis on Bray-Curtis values. The dominant explanatory variable was environment of origin (i.e., whether a sample came from the lab or field; NMDS mean stress ≈ 0.15, PERMANOVA, R^2^ = 0.150, Benjamini-Hochberg corrected p ≤ 0.001; Beta-dispersion F = 126.8, p ≤ 0.001; Table S1, Fig. 1). The R^2^ values (see Table S1) indicated that environment of origin was clearly an overarching explanatory variable, thus we decided to further analyse the field and lab samples separately to establish the important deterministic factors within each environment. The main trend in our results was a significant reduction in microbiome richness in lab-reared flies of all species based on a paired Wilcoxon rank sum test between Shannon index values for lab and field samples (B-H corrected p ≤ 0.001; Fig. 2). This significant trend held when pupae or adult *Drosophila* were analysed separately (Fig. S3, S4, S5).

**Figure 1:**
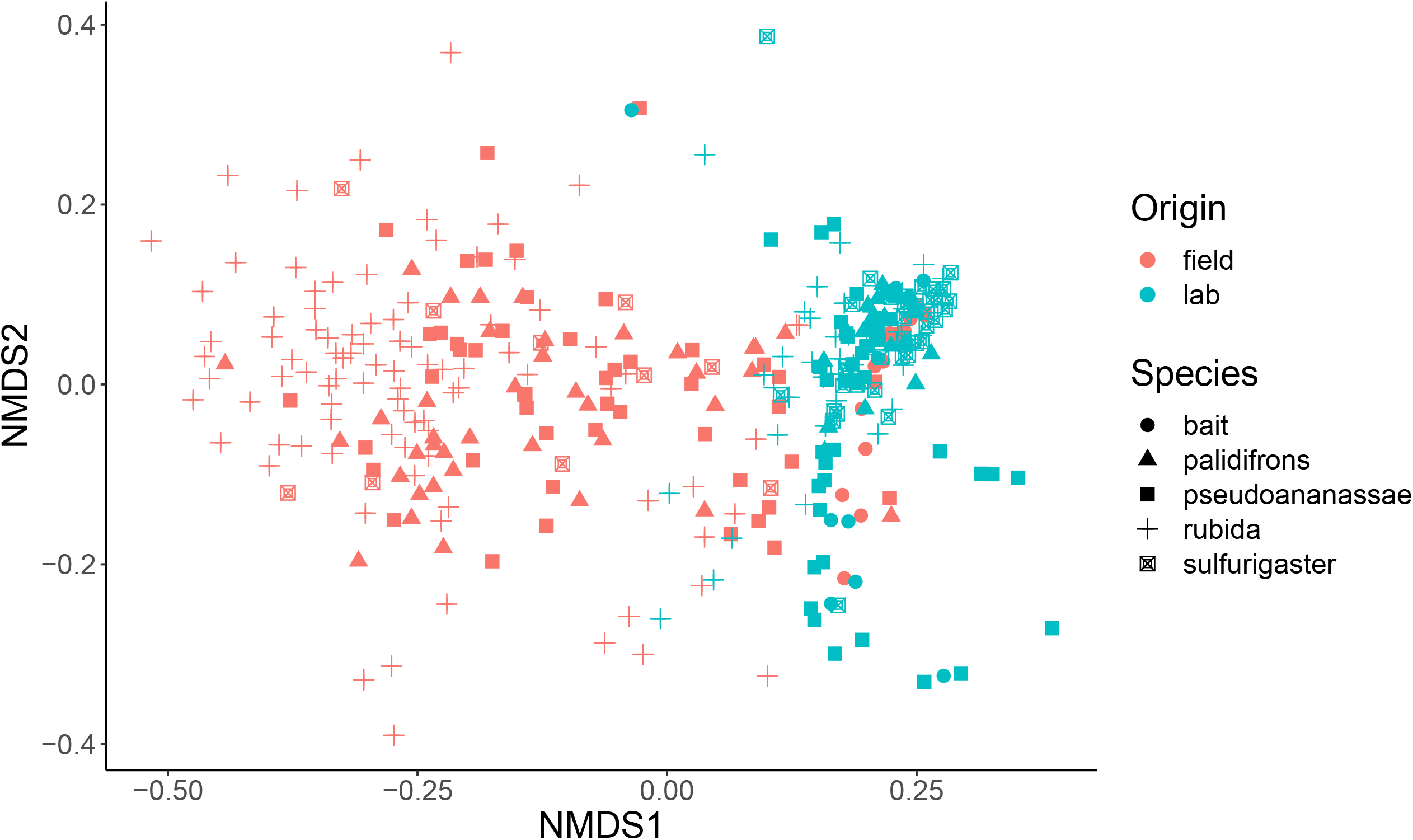
NMDS analysis of microbiome communities from all samples in this study, from the lab (blue) and the field (red). Samples of *Drosophila rubida, D. pseudoananassae, D. pallidifron*s, *D. sulfurigaster*, and food bait are indicated by different shapes.

**Figure 2:**
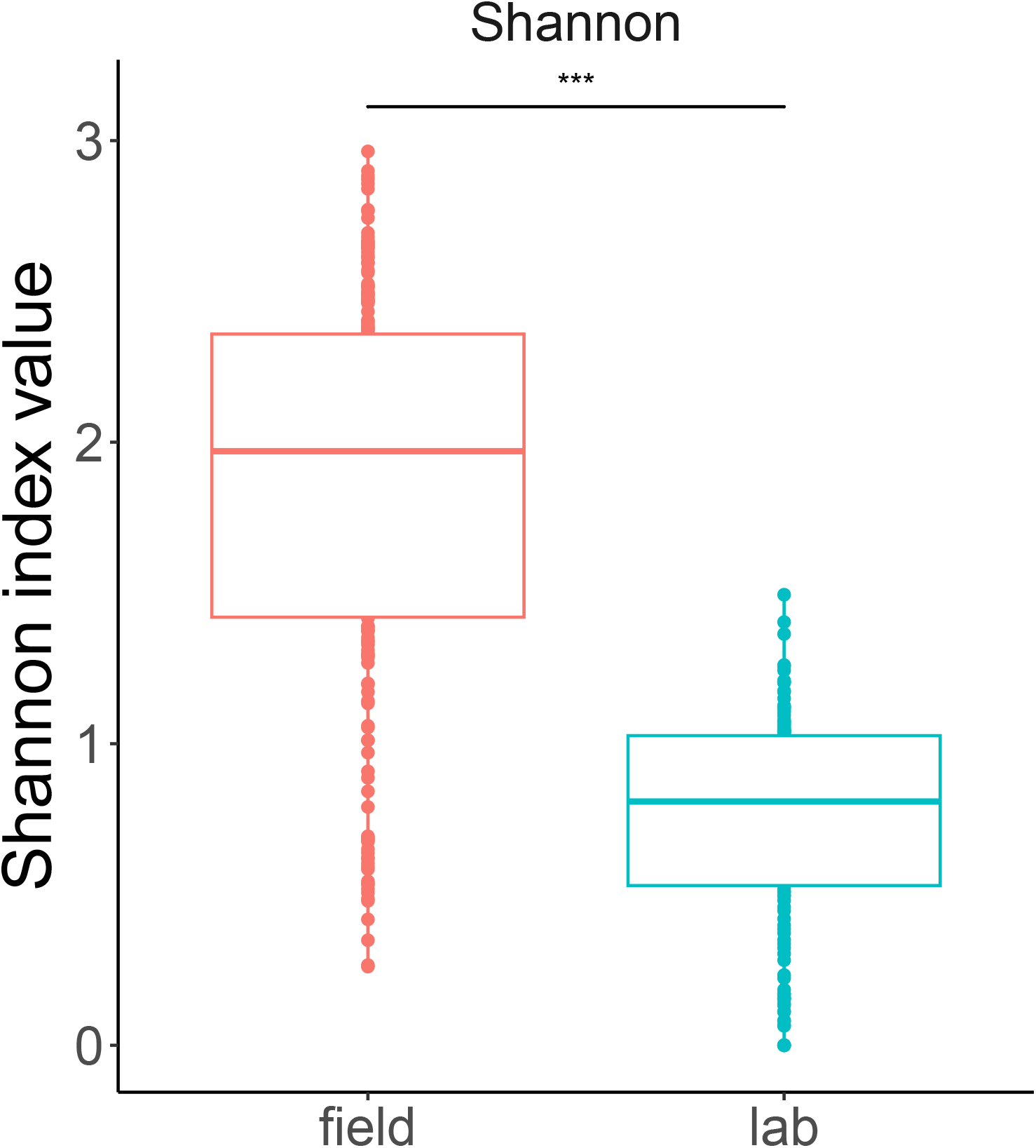
Comparison of Shannon index values for all samples in this study, split by environment of origin. Lab samples are shown in blue and field samples in red. The triple asterisk (***) shows high significance (p > 0.001) of a paired Wilcoxon test.

### 3.1 Microbiome patterns from field samples

When focusing on core field samples, the main factors explaining variation in microbiome structure were trap identity (PERMANOVA R^2^ = 0.13, B-H corrected p ≤ 0.001; Beta-dispersion p ≤ 0.001) and the interaction between site & trap (PERMANOVA R^2^ = 0.13, B-H corrected p ≤ 0.001; Beta-dispersion p ≤ 0.001; Table S2), suggesting that local environmental differences in site location, as well as trap identity, have a significant effect on *Drosophila* microbiome composition (Fig. 3). Elevation alone explained a small, but still significant, proportion of the variation observed (R^2^ = 0.03). The banana-baited bottle traps were left exposed in the field for different durations (between 11 and 24 days) to ensure we characterized the *Drosophila* community fully. PERMANOVA results suggest that the length of exposure had significant influence on microbiome composition but of lower importance compared to the other factors we identified based on the amount of variation explained (R^2^ = 0.03). Based on the PERMANOVA analysis, field site was a significant variable but only explained 7% of the variation in diversity (reflected in the minimal differences in average Shannon index value in Fig. S6). We also found evidence of *Drosophila* species-specific differences in microbiome diversity, based on paired Wilcoxon tests between Shannon index values for different species (p ≤ 0.001; Fig. 4). DeSEQ analysis of differential abundance indicated some bacterial genera were significantly more abundant in some *Drosophila* species but not others, most prominently *Kozakia* and *Corynebacterium* (Table S3). The most abundant bacterial genera (Fig. 5) were evenly distributed throughout all four *Drosophila* species sampled here, including *Acinetobacter*, which was the most dominant genus overall.

**Figure 3:**
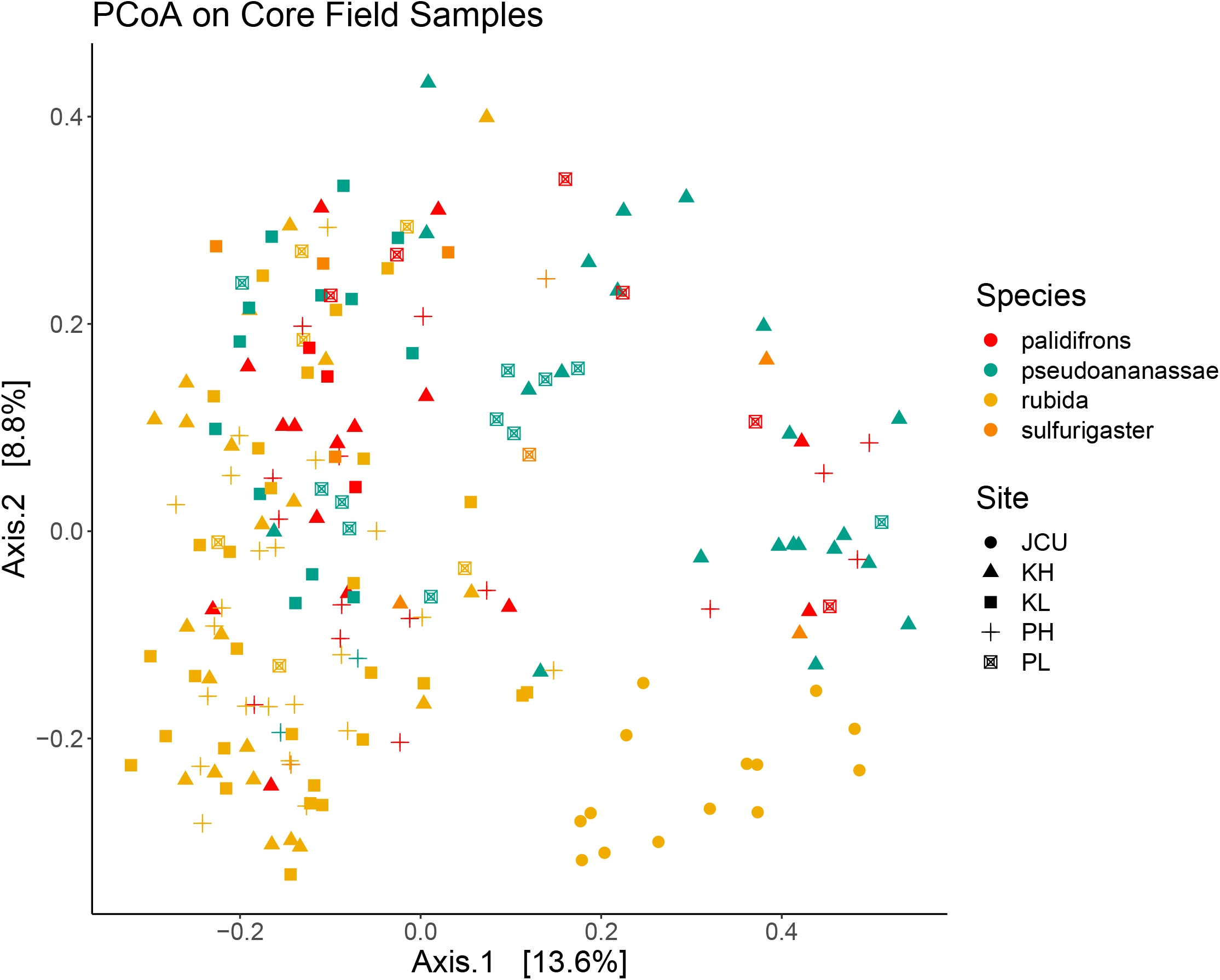
PCoA analysis of microbiome communities from the core field samples. Samples of *Drosophila rubida, D. pseudoananassae, D. pallidifron*s, and *D. sulfurigaster* are indicated by different colours, and the different sites of collection are indicated by shape. K = Kirrama, P = Paluma, JCU = James Cook University, L = Low [elevation], H = High [elevation]. Samples of *D. rubida* collected at JCU were all adults.

**Figure 4:**
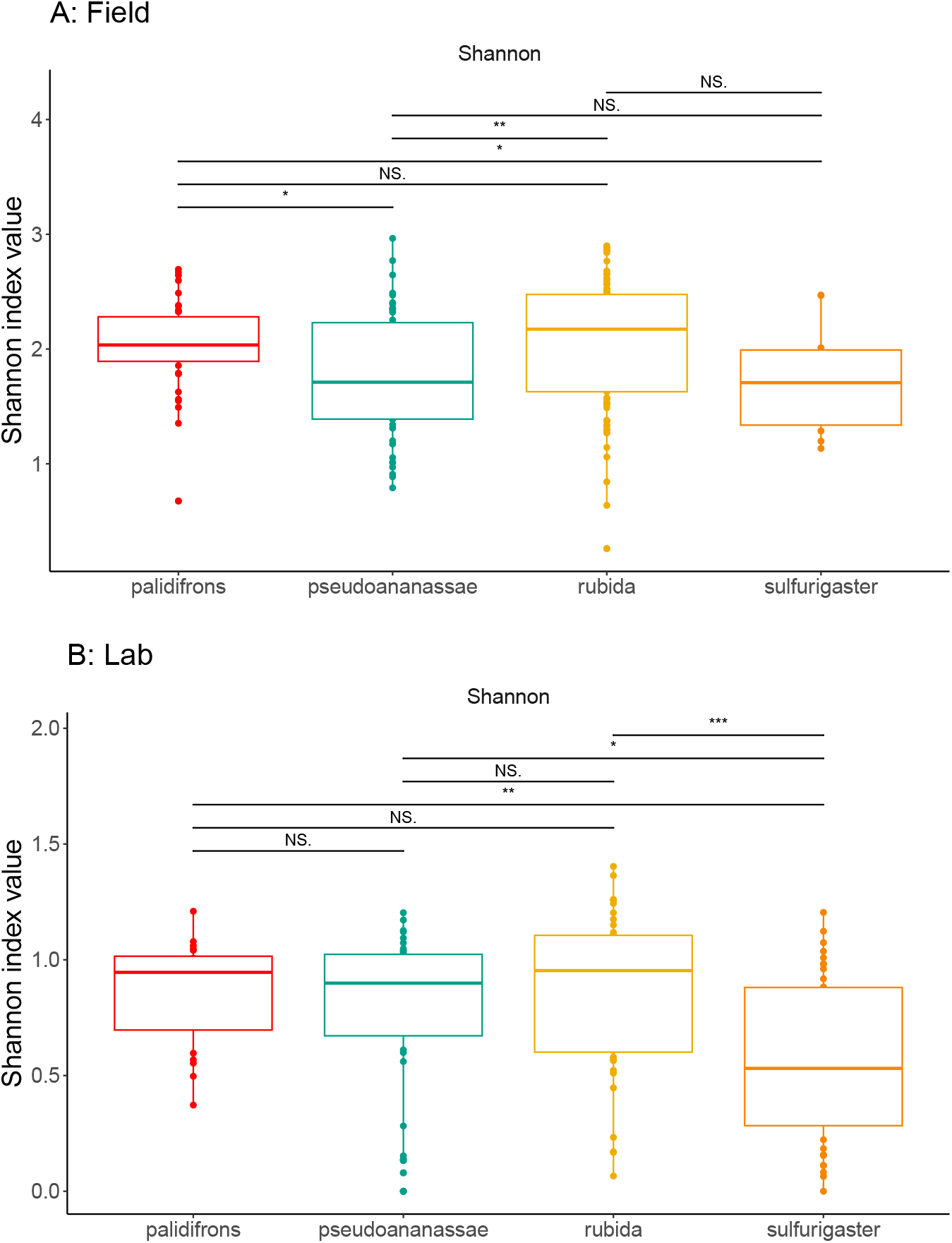
Comparison of Shannon index values from each species of *Drosophila* from (A) field and (B) lab environments. Each colour represents a different species. Statistical comparisons come from paired Wilcoxon tests. Three asterisks (***) denotes a highly significant result (p ≤ 0.001). Two asterisks (**) indicates a result of moderate significance (between p < 0.1 and p ≤ 0.001). One asterisk (*) denotes a marginally significant result (p < 0.1). NS. = not significant.

**Figure 5:**
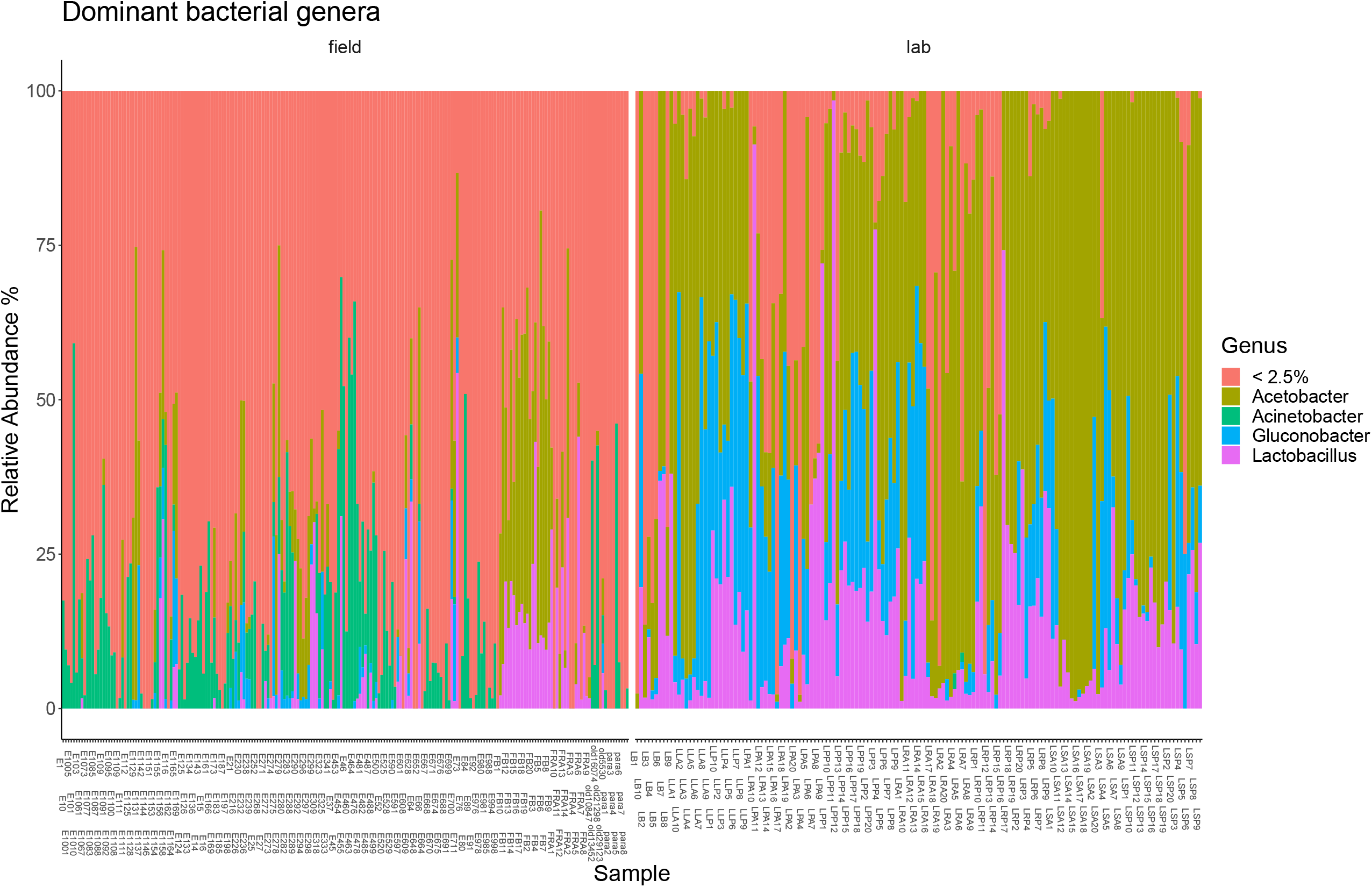
The most abundant bacterial genera from all samples in this study. Different colours mark different bacterial genera. <2.5% is a conglomerate category of low abundance taxa that made up less than 2.5% of the median number of reads. Each individual column represents an individual sample. Relative abundance is on the y axis.

### 3.2 Microbiome patterns from laboratory samples

The main factors explaining variation in microbiome structure from the lab-reared samples were *Drosophila* species identity (PERMANOVA R^2^ = 0.11, B-H corrected p ≤ 0.001, Beta-dispersion p ≤ 0.001), isofemale line (PERMANOVA R^2^ = 0.13, B-H corrected p ≤ 0.001, Beta-dispersion p ≤ 0.001), and the number of generations an isofemale line had been in the lab (PERMANOVA R^2^ = 0.12, B-H corrected p ≤ 0.001, Beta-dispersion p = 0.007; Table S4), suggesting that the duration a lineage of flies had been in the lab environment was important for explaining microbiome composition. The amount of variation explained by species-specific differences can largely be attributed to the presence of *Wolbachia* in *D. pseudoananassae* (Fig. 4 & Fig. S7), because the alpha diversity differences between species were not significant (Shannon index values, Fig. 6). All four *Drosophila* species contained high proportions of *Acetobacter, Lactobacillus* and *Gluconobacter*, but only *D. pseudoananassae* contained *Wolbachia*. The interaction between isofemale line and generation was also highly significant and explained a high proportion of variation relative to other factors, which was expected because IF lines were established at different times, and therefore had been in the lab for varying numbers of generations at the time of sampling. Life stage (i.e., the difference between pupae and adults) was not one of the most important explanatory factors based on the amount of variation explained (PERMANOVA R^2^ = 0.06). There was no detectable effect of elevation (PERMANOVA R^2^ = 0.01), i.e., the elevation of the field site where an isofemale line was collected did not have a significant effect on the microbiome structure in the lab.

**Figure 6:**
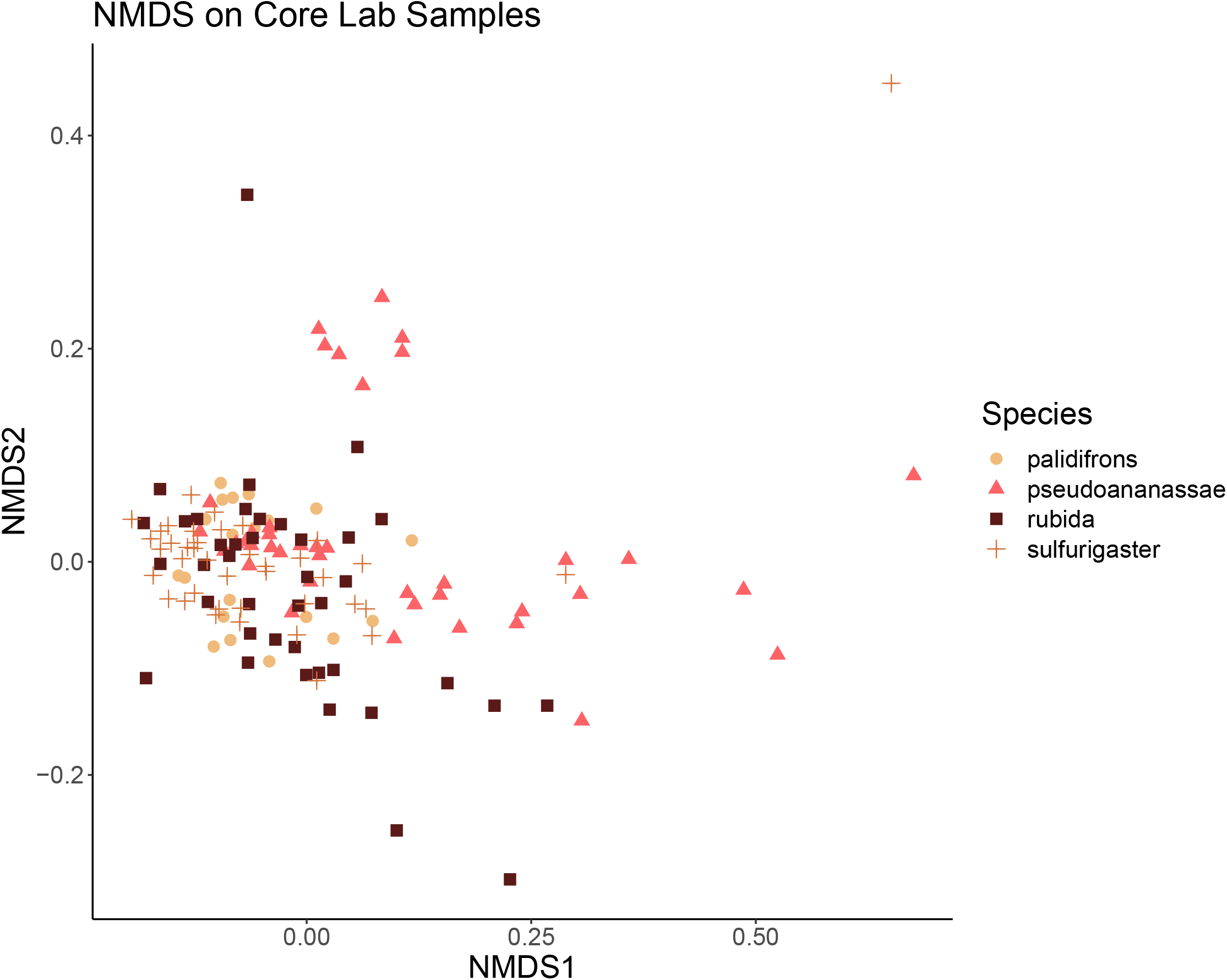
NMDS analysis of microbiome communities from core lab samples. Samples of *Drosophila rubida, D. pseudoananassae, D. pallidifrons*, and *D. sulfurigaster* are indicated by different colours and shapes.

### 3.3 Comparisons between field and laboratory samples and microbiome of the food source

There was no significant difference in microbiome composition of laboratory food samples from Czech Republic and the banana bait that we used in the field in Australia (Paired Wilcoxon test on Shannon index values, p = 0.09; Fig. S8). Banana bait and lab fly food both contained high relative proportions of *Acetobacter* and *Lactobacillus*, with some lab food samples containing *Gluconobacter* (Fig. S9). Despite the physical differences in food sources, the most abundant bacterial genera were the same, and the food source microbiome diversity was low. In lab-reared flies, these three bacterial genera dominated the microbiomes of pupae and adults (Fig. S7). In the field, however, *Acetobacter* and *Lactobacillus* were not the most abundant genera of either pupae or adult gut microbiomes. *Acetobacter* and *Lactobacillus* had the greatest difference in relative abundance between field and lab samples (summary statistics in Table 2, Fig. S10). Whilst still present in field samples, the relative abundance of *Acetobacter* and *Lactobacillus* was proportionally lower in their taxa-rich microbiomes (Fig. 1).

**Table 2:** Summary output of DeSEQ differential abundance analysis showing the top 5 bacterial genera by origin of sample. Positive test statistic values indicate a genus significantly more abundant in lab samples and negative test statistics indicate a genus significantly more abundant in field samples.

| Genus | baseMean | log2foldchange | lfcSE | Test statistic | Adjusted p value |
| --- | --- | --- | --- | --- | --- |
| ***Acetobacter*** | 436.99 | 3.59 | 0.38 | 9.46 | ≤ 0.001 |
| ***Lactobacillus*** | 183.75 | 3.16 | 0.38 | 8.34 | ≤ 0.001 |
| ***Acinetobacter*** | 195.62 | -7.91 | 0.33 | -23.70 | ≤ 0.001 |
| ***Klebsiella*** | 105.22 | -7.01 | 0.37 | -18.99 | ≤ 0.001 |
| ***Chishuiella*** | 59.69 | -6.19 | 0.41 | -14.97 | ≤ 0.001 |

### 3.4 Robustness of results

There was no detectable difference in microbiome diversity of parasitised pupae of *D. rubida*, compared to unparasitized pupae (Paired Wilcoxon test on Shannon index values, p = 0.25; Fig. S11), nor were there bacterial genera unique to parasitised samples. We also found that the 99% identity OTU table produced qualitatively the same results as the 97% identity OTU table. For example, in an NMDS on all samples the dominant explanatory variable was still environment of origin (NMDS mean stress ≈ 0.17, PERMANOVA, R^2^ = 0.17, B-H corrected p ≤ 0.001; Beta-dispersion p ≤ 0.001; Fig. S1). Removing *Wolbachia* from the dataset also did not qualitatively change the outcome of statistical tests, e.g., the main factors explaining microbiome structure in lab-reared samples were *Drosophila* species identity (PERMANOVA R^2^ = 0.11, B-H corrected p ≤ 0.001, Beta-dispersion p ≤ 0.001), isofemale line (PERMANOVA R^2^ = 0.17, B-H corrected p ≤ 0.001, Beta-dispersion p ≤ 0.001), and the number of generations an isofemale line had been in the lab (PERMANOVA R^2^ = 0.12, Benjamini-Hochberg corrected p ≤ 0.001, Beta-dispersion p ≤ 0.001).

## 4. Discussion

Our results revealed small but significant variation in microbiome structure between *Drosophila* populations from high and low elevation across both gradients. We expected to find greater differences across elevation, because there is well-documented evidence of both insects and bacteria developing differently according to differences in temperature of >5°C^29–31,33,34,64^. This finding could be a result of the *Drosophila* species sampled here being ubiquitous across elevation without forming sufficiently distinct populations at high and low elevation sites, or because the ∼5°C temperature shift between our sites was not strong enough to drastically alter microbiome composition. Standardising diet may have homogenised *Drosophila* microbiomes to some extent, but the overall lack of similarity in microbiome composition between food microbiome and field-caught pupae suggests that diet was not a dominant variable in structuring *Drosophila* microbiomes.

The most pronounced differences in microbiome composition were between individuals raised in the laboratory and those raised in the field. Interestingly, the two food sources (banana bait in the field and yeast-based *Drosophila* medium in the lab) had very similar microbiome profiles, suggesting that dietary factors were not primarily responsible for the observed differences between environments. The bacterial community from lab food matches well with the microbiomes found within lab-reared pupae and adult flies. This was expected, because it reflects a well-established pathway of insect microbiome colonisation - *Drosophila* ingest food and acquire bacteria associated with that food source^36^. Yet in the field, *Drosophila* microbiomes do not correspond well with the bacterial communities found on banana bait samples. The observed pattern can be explained by significant differences in microbiome colonisation from environmental bacterial species pools^65,66^. The flies sampled from the lab come from a highly regulated environment, with a specific and consistent food source provided into heat-sterilised glass vials, so the only ‘available’ bacteria for colonising their microbiomes come from the diet, surrounding lab environment, and vertically inherited endosymbionts (e.g., *Wolbachia* in *D. pseudoananassae*). In contrast, the bacterial species pool in Australian tropical rainforest comprises much greater diversity and abundance of different taxa, creating a greater variety of possible microbiome composition within *Drosophila* hosts. This diversity of taxa creates more room for ecological drift, dispersal, and selection to act on microbiome communities, in turn creating greater among-individual and between-species variation in wild flies. Consistently higher diversity in wild *Drosophila* microbiomes suggests that microbiomes are predominantly colonised from the wider environment and dependent on local species pool diversity. For instance, bottle traps were visited by other organisms - e.g., staphylinid beetles, neriid flies, Lepidoptera - which could also have been a source of bacteria colonising the microbiome of the *Drosophila* sampled in this study.

There was consistently low microbiome diversity found in both lab-reared pupae and lab-reared adults, suggesting that low diversity within pupae is an accurate representation of lab-reared microbiomes. This result was surprising because we anticipated some stage-specific microbiome community patterns, given that *Drosophila* are holometabolous and thus undergo substantial gut remodelling during complete metamorphosis^67^. The consistency across life stages from lab-reared individuals provides further evidence for the depauperate nature of the lab microbial environment. In contrast to the lab, the field-caught adults of *D. rubida* lacked congruence with the field-caught pupae, and both stages lack similarity with the banana bait (Fig. S12, S13). The lack of geographic variation in microbiome composition suggests that different sites are unlikely to fully explain the discrepancy. With adult flies we cannot rule out that they might have fed on a substance other than our yeasted banana bait prior arriving at our bottle traps, making dietary variation a parsimonious explanatory factor in life-stage differences^36,68^. It is also possible that the high diversity we found in pupal samples reflects their metabolic activity (despite their lack of feeding), given that complete metamorphosis is an intense period of organismal change^69^. The substantial differences in microbiomes between lab and field specimens suggests that future studies should be cautious and specific with the types of microbiome-related questions investigating laboratory flies. Interpreting microbiome community composition from lab-kept specimens, particularly those from cultures maintained across multiple generations, is unlikely to yield data entirely representative of natural microbiomes^70,71^, except perhaps in scenarios where microbiome communities from lab-reared specimens are a subset of the more diverse microbiomes found in the field (as our results show in Figure 5).

Previous studies on *Drosophila* have demonstrated high intra- and interspecific variation in microbiome community composition from both wild-caught and lab-reared flies^39,40,70,72,73^, which our results corroborate. Controlling for diet in both scenarios allowed us to recognise this species-specificity more accurately. We found a significant effect of species identity (11% variation explained) but it did not explain as much variation as the results from Adair et al.^40^ who found species identity explaining 42% and 70% variation in two different sets of *Drosophila spp*. This discrepancy could be a product of the species themselves (i.e., we used a different set of *Drosophila* species) or the number of species studied (we studied four species here; Adair et al.^40^ eighteen), but the evidence from both studies suggests that species-specificity is maintained in the lab, albeit not in a consistent manner. Statistically significant differences in microbiome alpha diversity measures between different *Drosophila* species in the field were not reflected in the lab. From the field, the only significant difference in microbiome richness was between *D. pseudoananassae* and *D. rubida*, and this was non-significant in the lab. Moreover, there was a highly significant difference in richness between *D. rubida* and *D. sulfurigaster* in the lab, but not in the field. Some of our significant PERMANOVA results were accompanied by significant Beta-dispersion, which suggests that there was some heterogeneous dispersion in the variables we measured. This was expected based on the high variation typically exhibited by microbiome community data and typically found when sampling wild individuals. Even with greater and more even sample size, we would still expect to find a large, significant difference in community richness and composition between field and lab *Drosophila* microbiomes.

In other insect species, host transmission of extracellular symbionts (like those in the gut) have been hypothesised to result in long-term associations between insect and microbe^74–76^. The long-term laboratory survival of our four *Drosophila* species (minimum 24 generations) with radically different microbiome composition (compared to their counterparts from the field) suggests that their symbiotic relationship with bacteria is not highly specialised^77^, corroborating previously published findings. For example, Wong et al.^37^ found no consistent evidence for a core microbiome in multiple *Drosophila* species and Storelli et al.^78^ described axenic *Drosophila* returning to normal growth and development in the presence of a single bacteria species. Furthermore, Coon et al.^79,80^ showed that any bacterium was sufficient to facilitate mosquito larval development, whereas axenic mosquito larvae do not develop past the first instar. Thus, our results provide further evidence of satisfactory microbiome function being provided by a limited number of bacterial species, which poses questions about the advantages of diverse microbiomes within wild insects and the links between microbiome community diversity and host fitness.

Overall, we found significant differences in microbiome diversity of field-caught and lab-reared *Drosophila*, which were consistent across species and life stage. We hypothesise that these differences in diversity are the products of environments with markedly different bacterial species pools. To elucidate functional conclusions from insect-microbiome analyses, more in-depth molecular analysis (e.g., metagenomics, transcriptomics) is required. We suggest that comparative studies between lab and field specimens help reveal the true variability in microbiome communities that can exist within a single species.

## Acknowledgements

We thank Hana Konvičková and Jan Zima Jr. for their assistance in the molecular lab. We acknowledge funding support from the Czech Science Foundation grant no. 17-27184Y to JH. OTL was further supported by the UK National Environment Research Council (NE/N010221/1). JH was additionally supported by the Czech Ministry of Education, Youth and Sports grant ERC CZ LL2001. Computational resources were supplied by the project “e-Infrastruktura CZ” (e-INFRA LM2018140) provided within the program Projects of Large Research, Development, and Innovations Infrastructures. Fieldwork was conducted under permit WITK16977516 from Queensland’s Department of Environment and Heritage Protection.

