## Supplemental material for "Microbiome structure of a wild *Drosophila* community along tropical elevational gradients and comparison to laboratory lines"

**Supplementary Information**

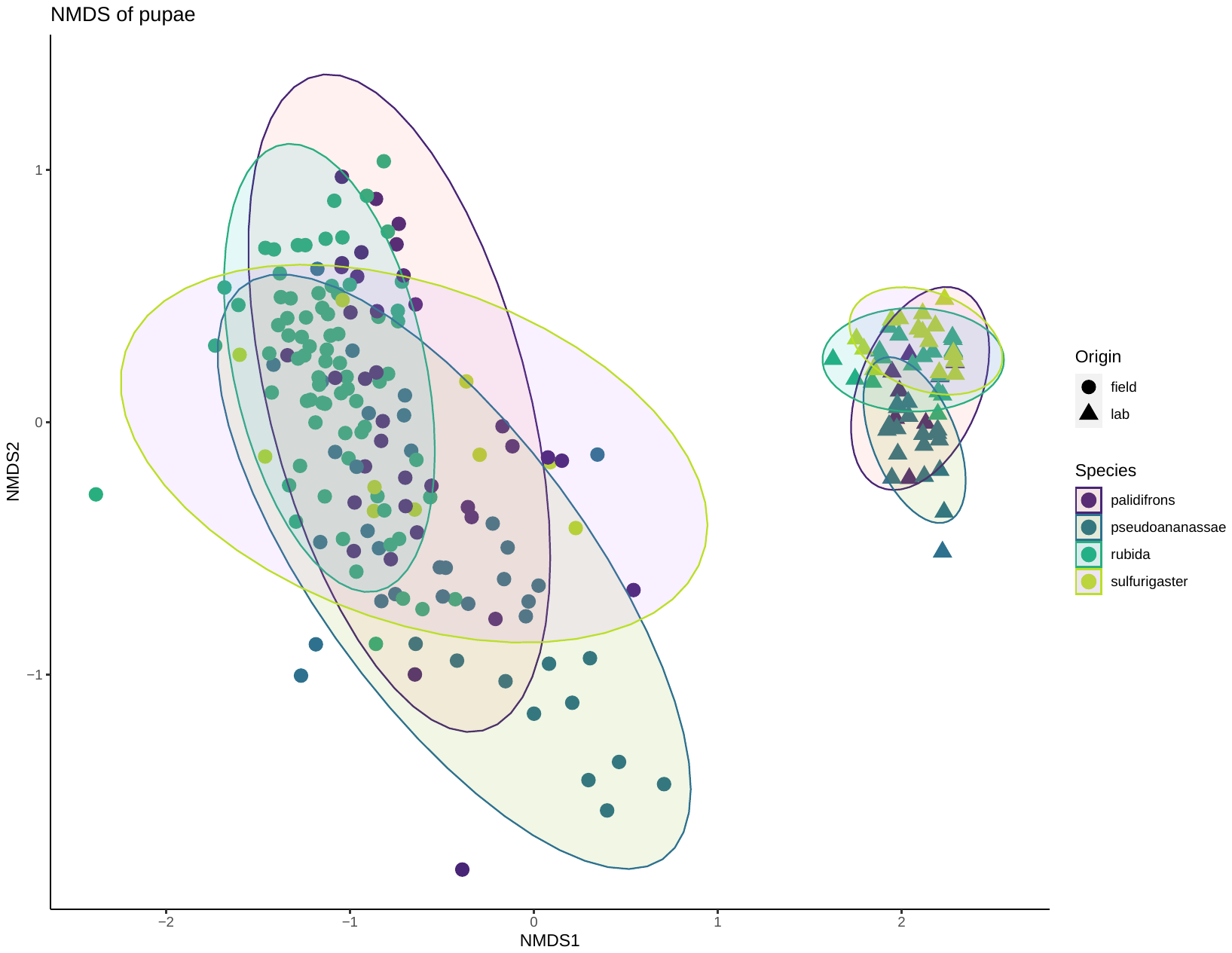

**Figure S1: NMDS of microbiome communities from all pupae in this study. Symbols represent site of origin and different colours represent different species. This result was generated from the dataset clustered at 99% sequence identity.**

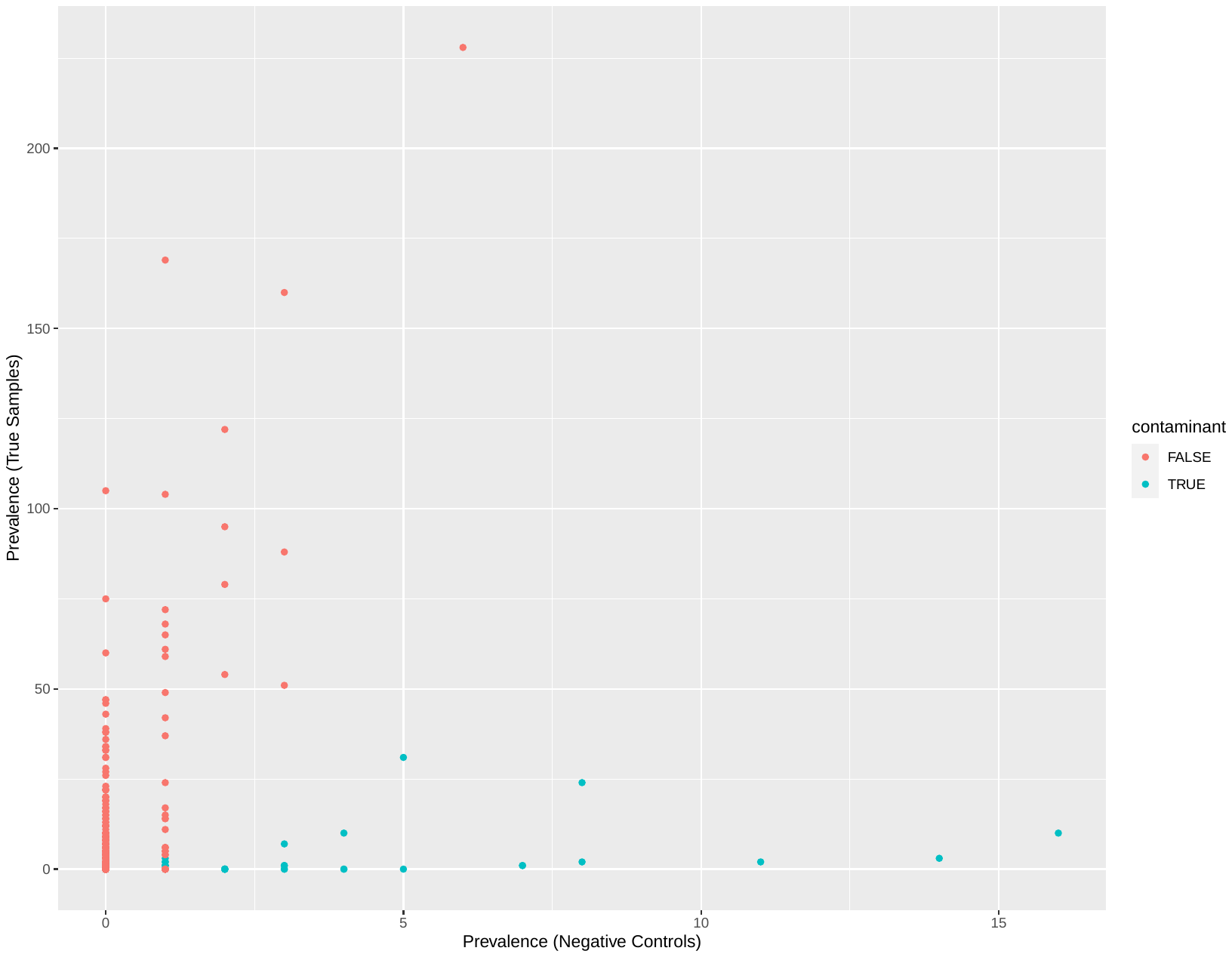

**Figure S2: Plot of contaminant OTUs identified by 'prevalence' function in the R package 'decontam'.**

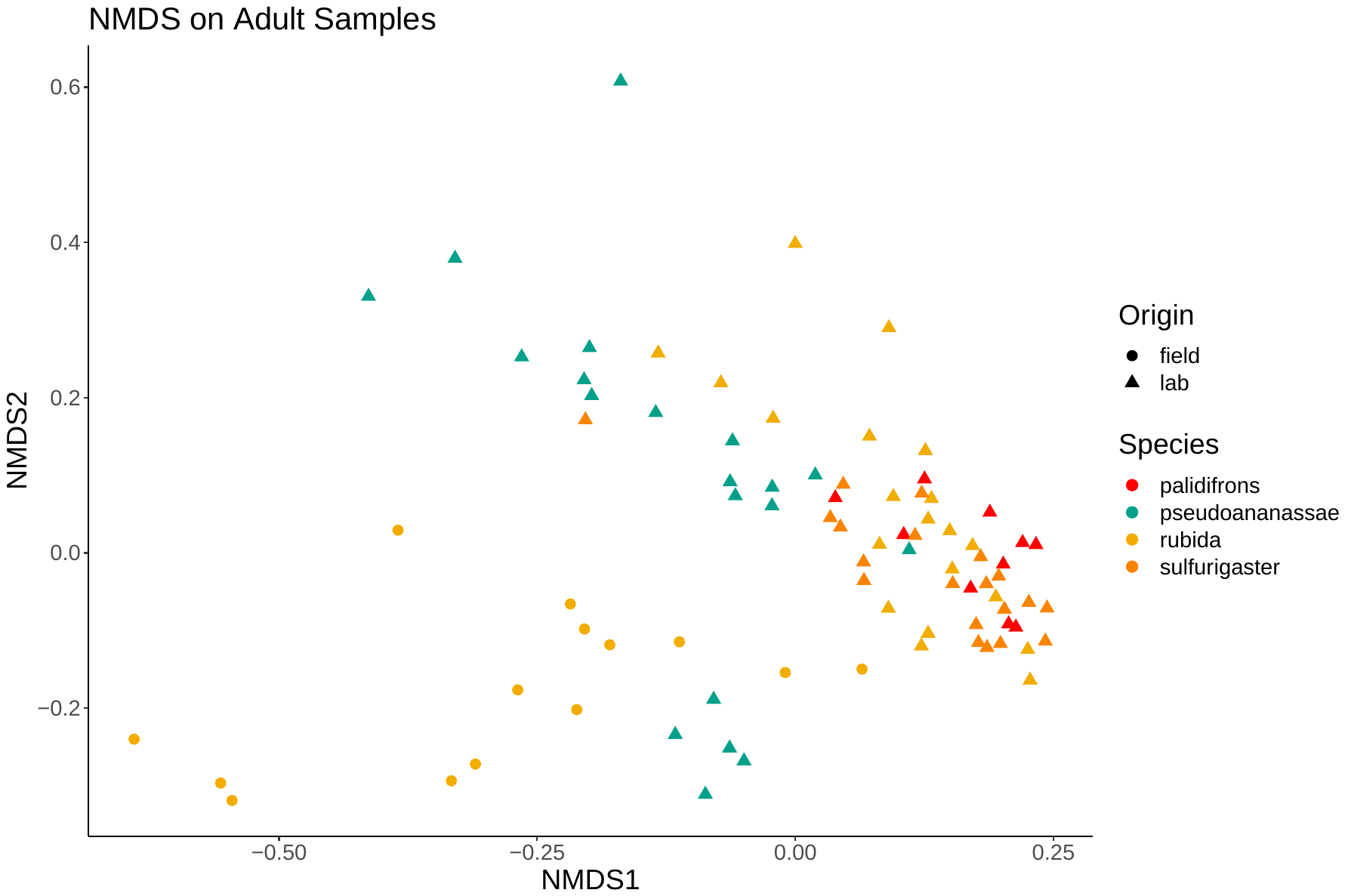

**Figure S3: NMDS of microbiome communities from all adults in this study. Symbols represent site of origin and different colours represent different species.**

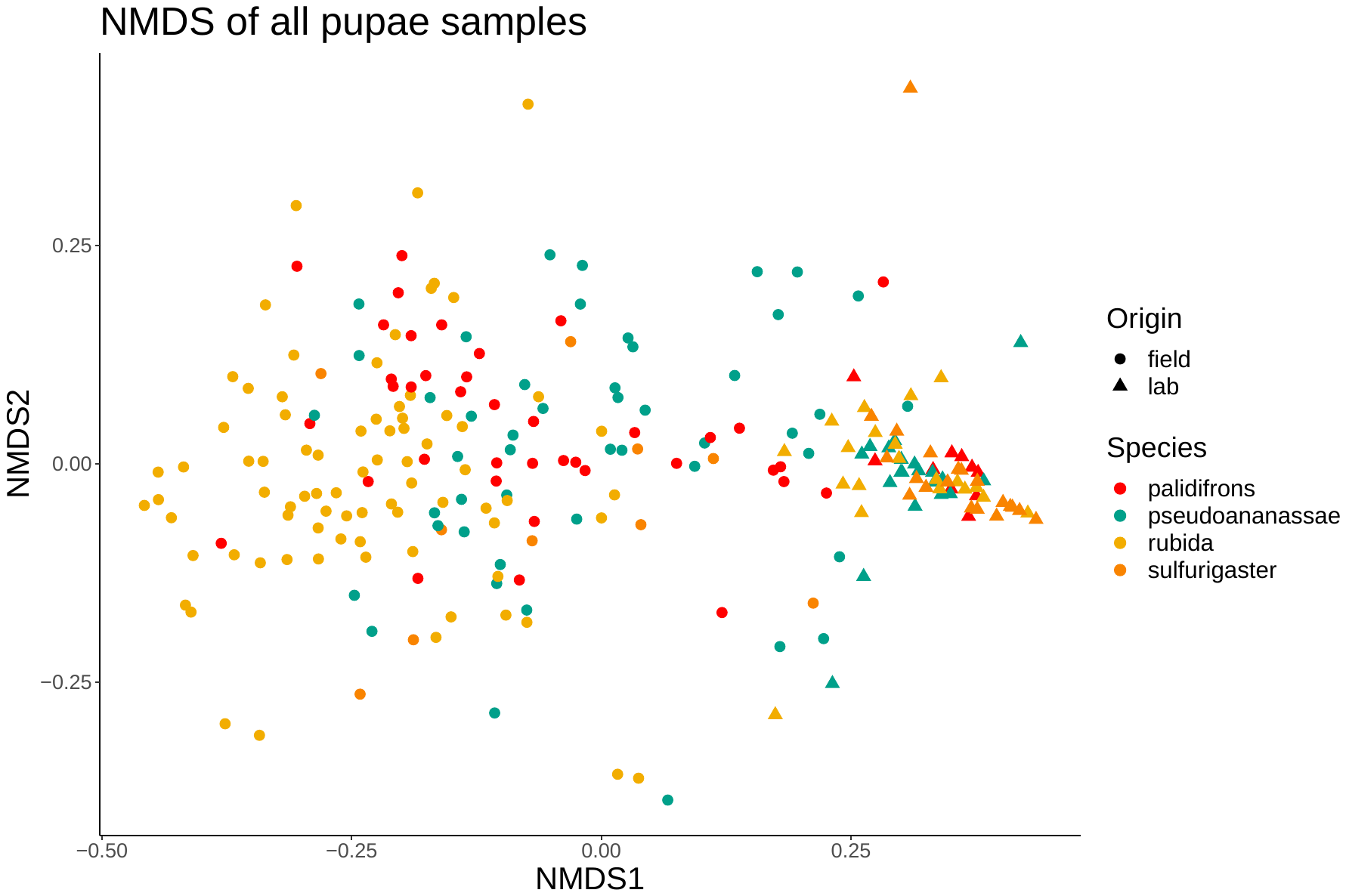

**Figure S4: NMDS of all pupae samples, showing the difference between environment of origin (circles vs triangles). Different colours represent different *Drosophila* species.**

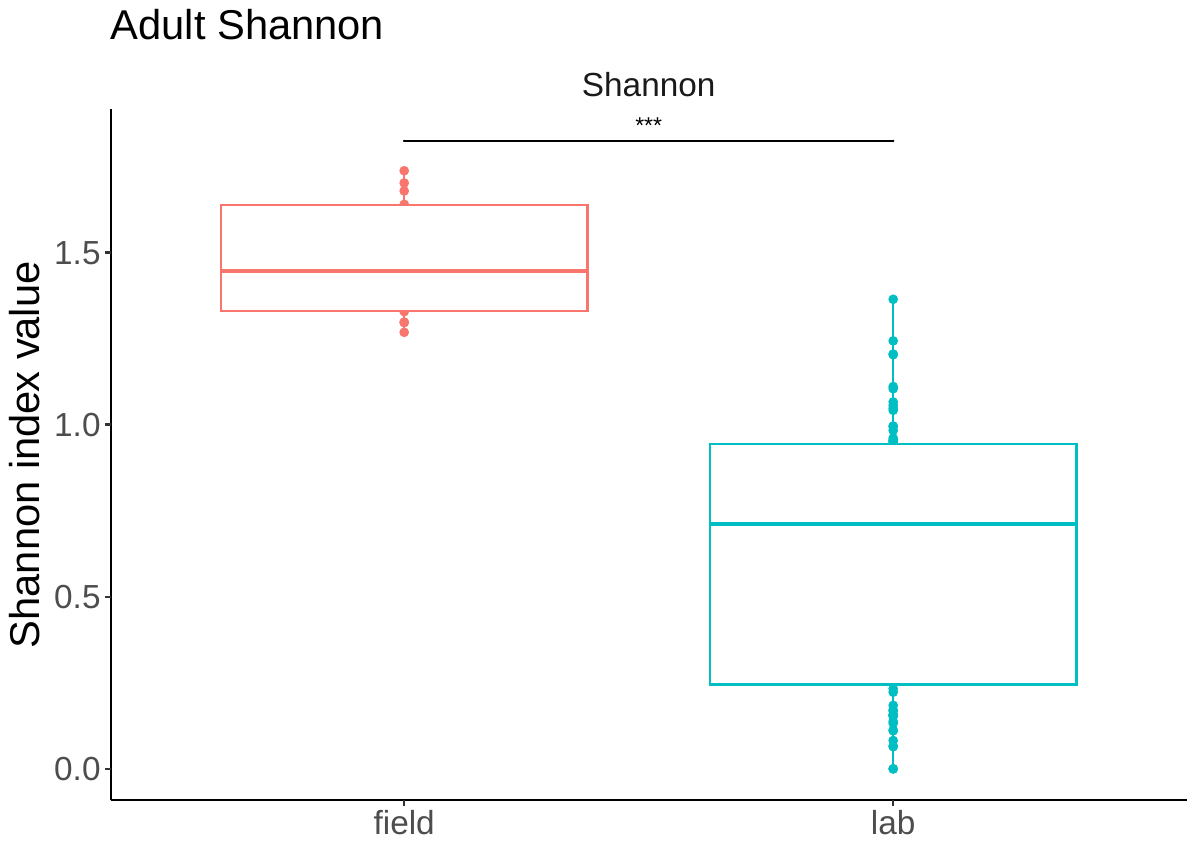

**Figure S5: Shannon index values for microbiome communities from all adults in this study. Colours denote environment of origin.** Three asterisks (***) denotes a highly significant result (p ≤ 0.001), based on paired Wilcoxon tests.

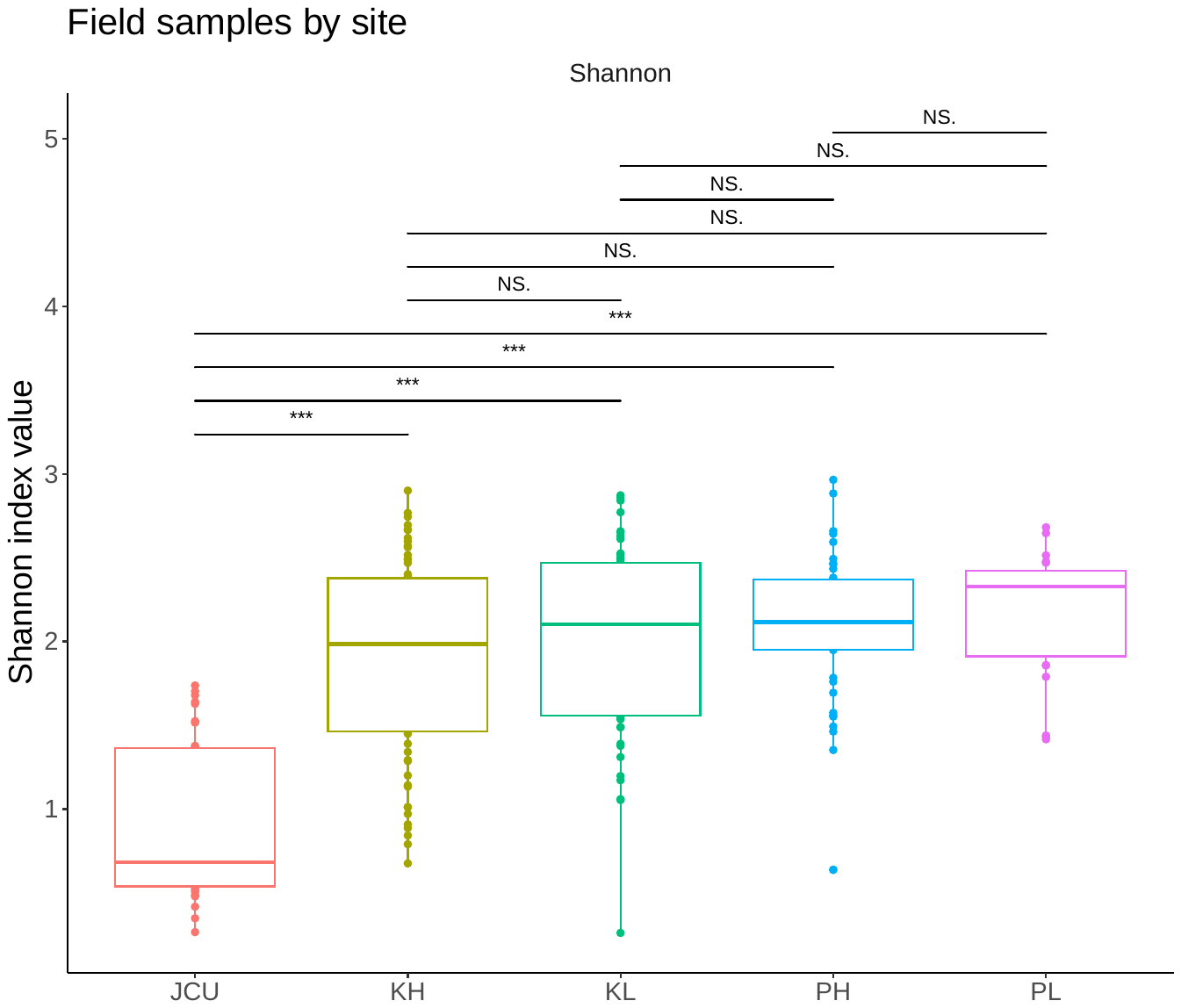

**Figure S6: Shannon index values for all field samples organised by site.** K = Kirrama, P = Paluma, JCU = James Cook University, L = Low [elevation], H = High [elevation].

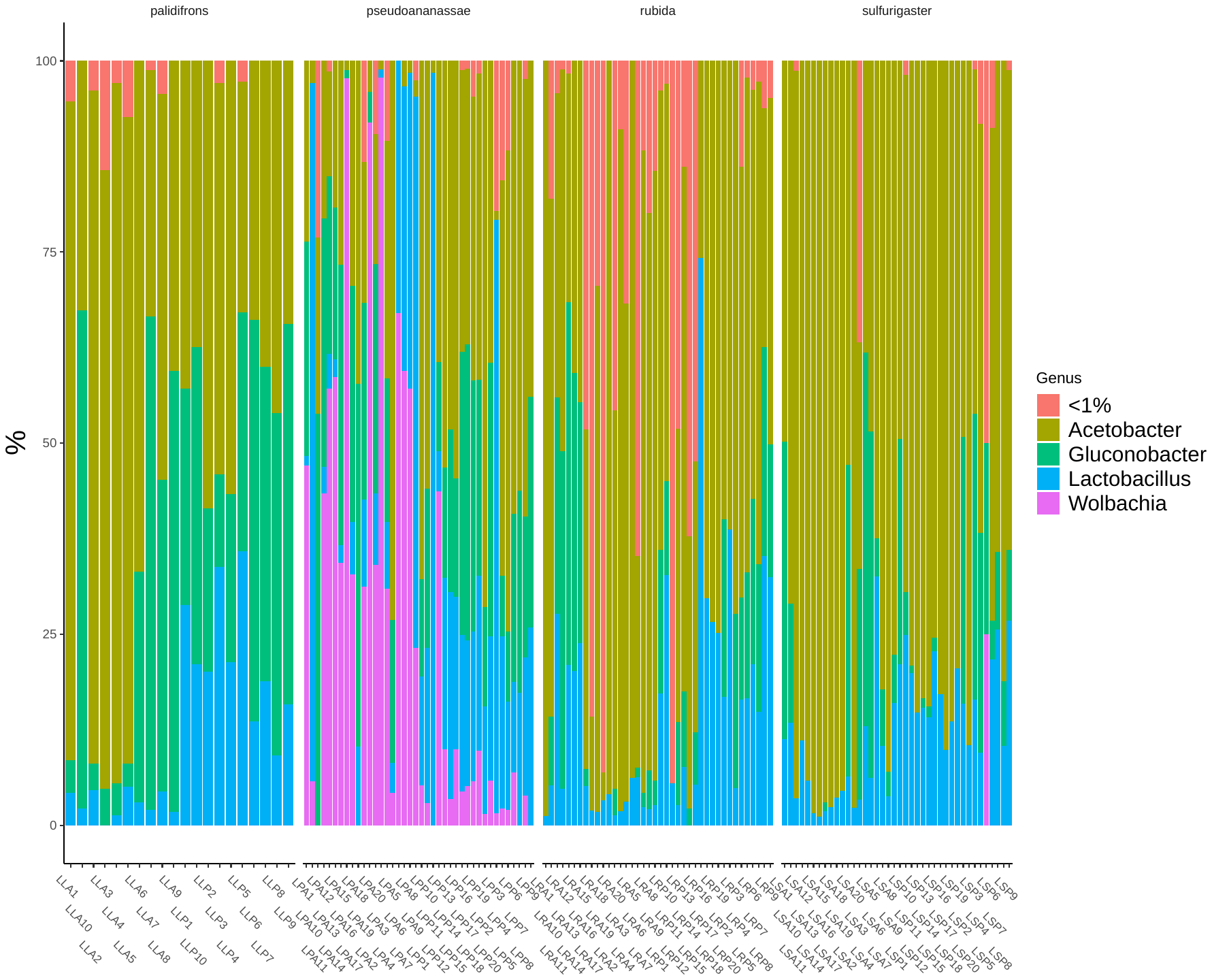

**Figure S7: The most abundant bacterial genera for lab-reared samples of all four *Drosophila* species.** Different colours represent different bacterial genera. Each individual column represents an individual sample. <1% is a conglomerate category of low abundance taxa that made up less than 1% of the median number of reads.

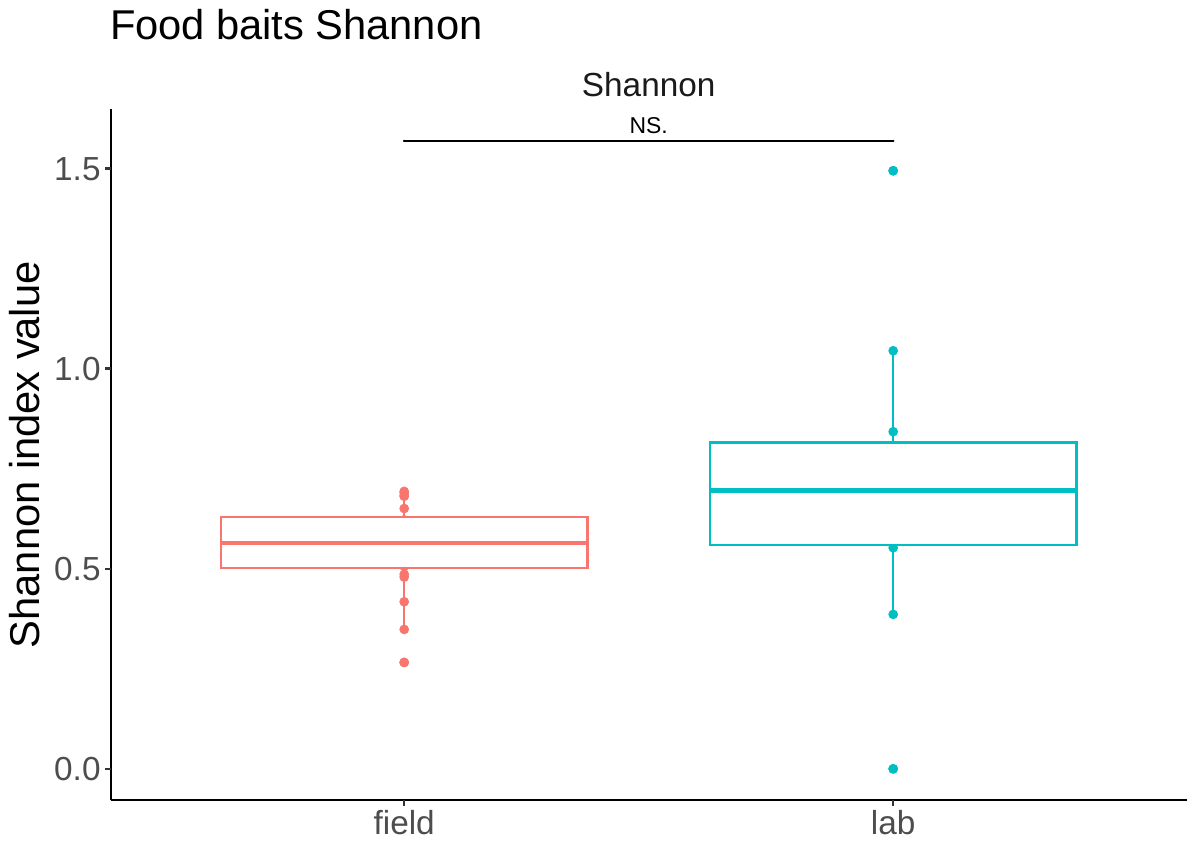

**Figure S8: Shannon index values for microbiome communities from bait samples in this study. Colours denote environment of origin.** NS. denotes a non-significant result (p > 0.05), based on paired Wilcoxon tests.

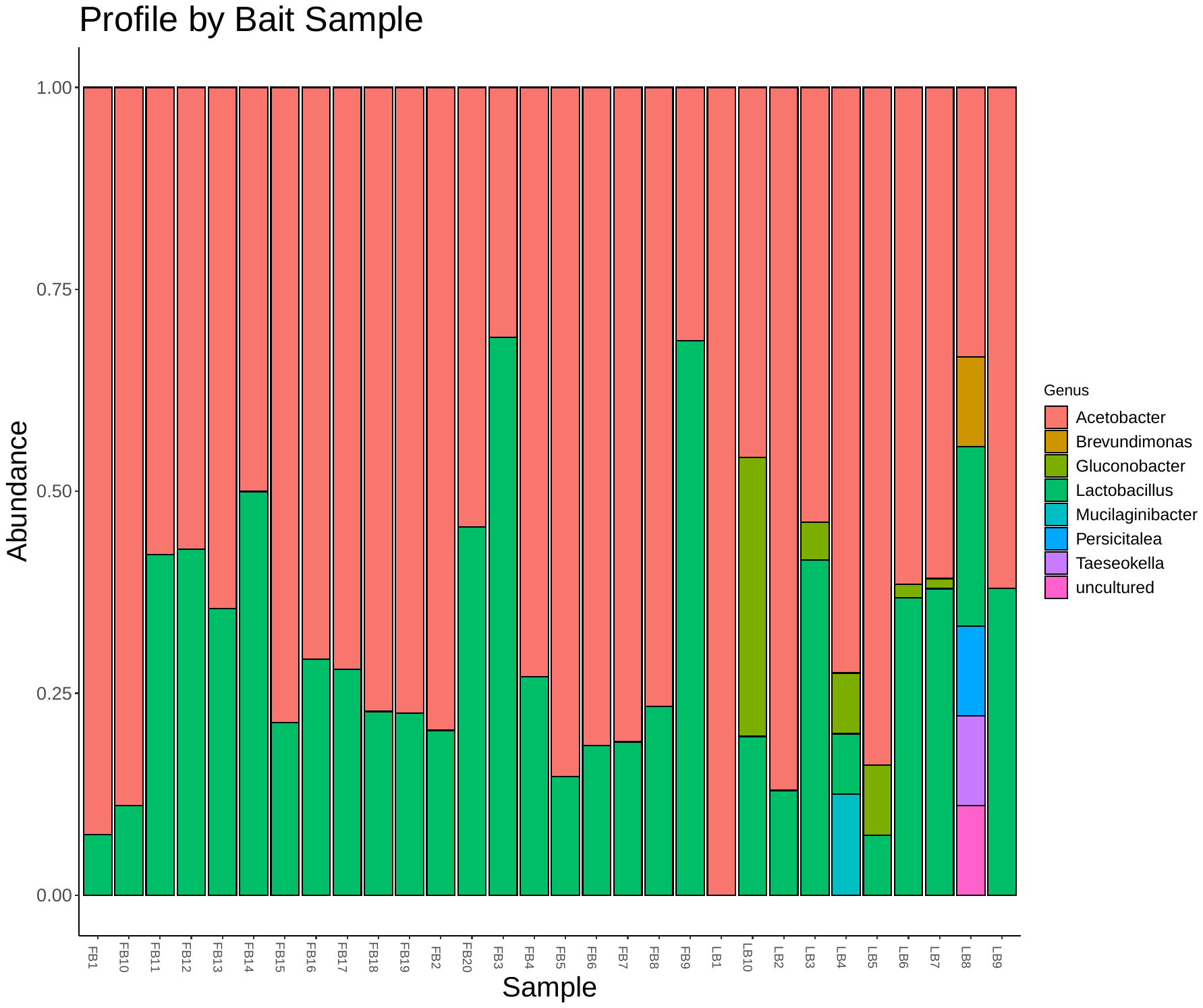

**Figure S9: Microbiome community profiles of the food baits used in this study.** 'Field' = banana bait used in bottle traps in Queensland, 'lab' = the mixture of yeast and agar used to feed *Drosophila* housed in laboratory conditions. y axis = relative abundance of each taxa.

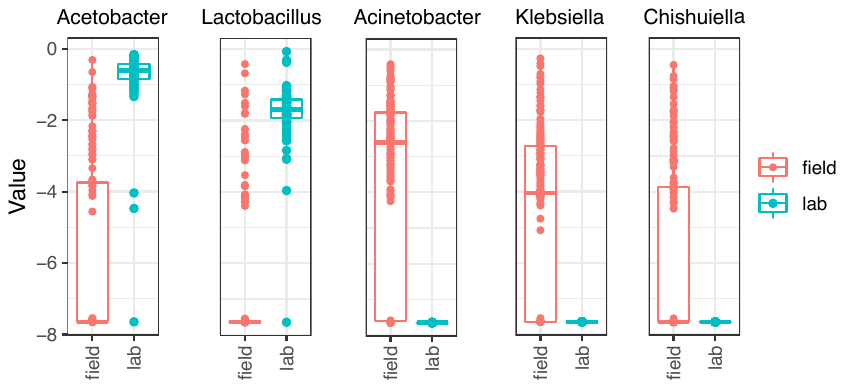

**Figure S10: DeSEQ analysis showing the most differentially abundant bacterial genera between lab and field core samples. y axis = DeSEQ test statistic difference.**

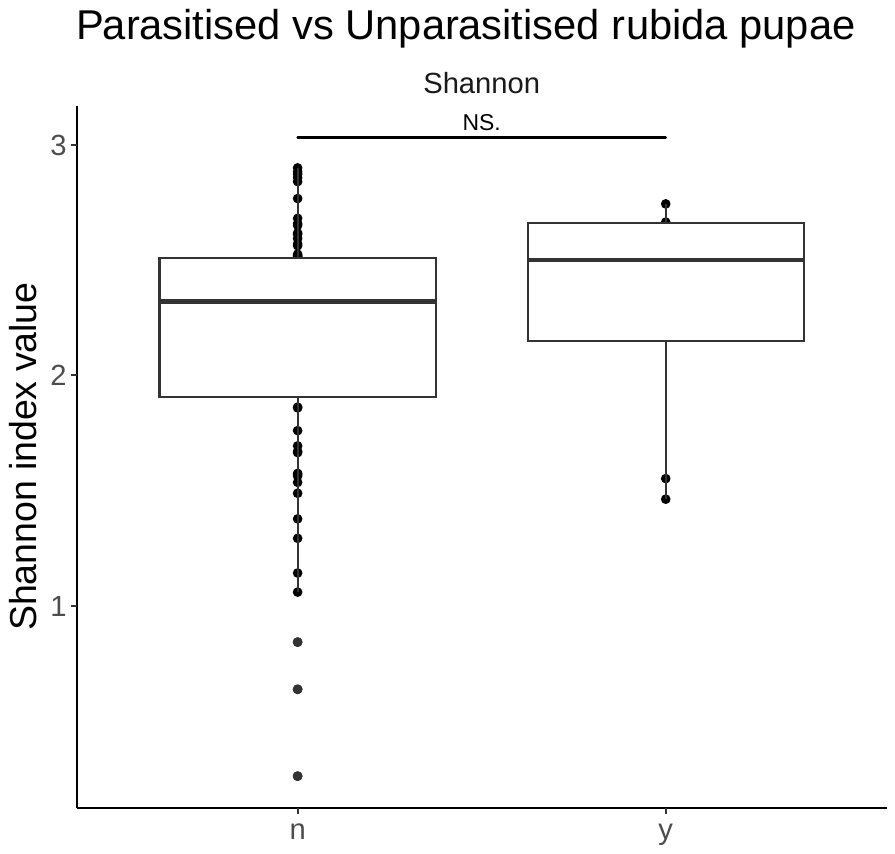

**Figure S11: Shannon index of all *Drosophila rubida* pupae sampled from the field, with the samples split by whether a sample had been parasitised by a parasitoid wasp.** N = no parasitoid, y = yes, positive detection of a parasitoid. NS. denotes a non-significant difference based on paired Wilcoxon tests.

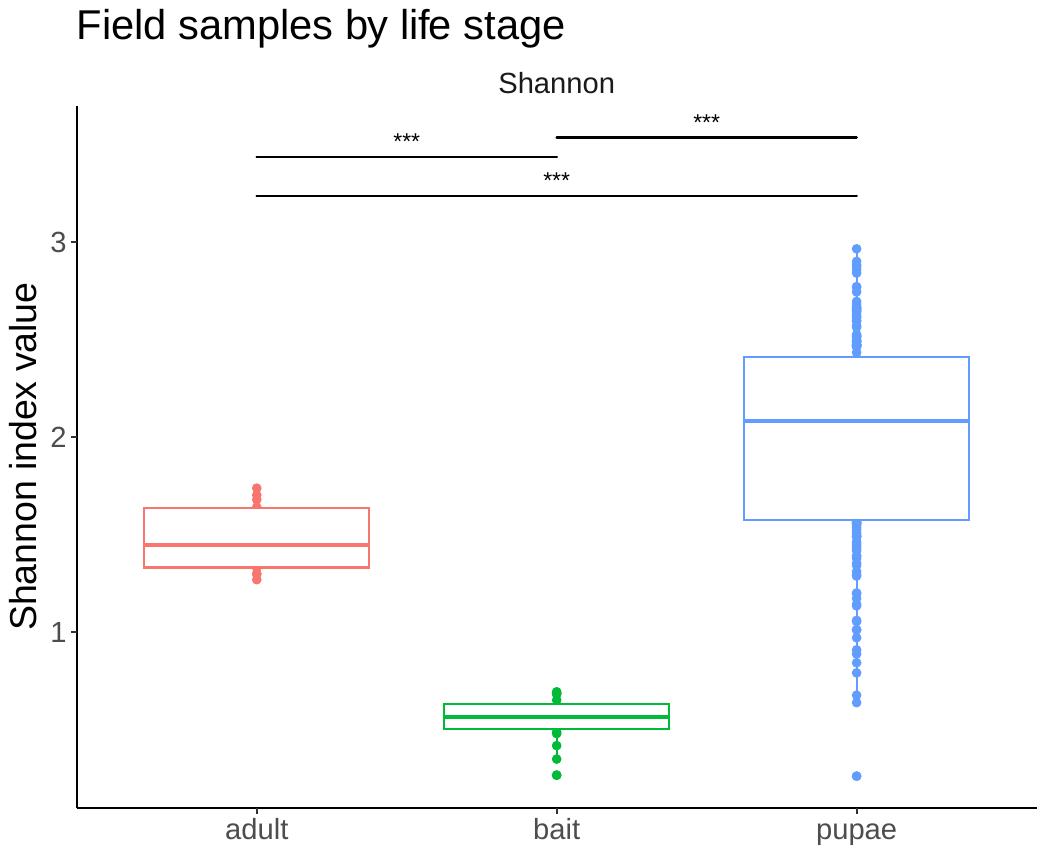

**Figure S12: Shannon index values for field bait and field core samples, split by life stage. Bait = banana baited fly food we collected *Drosophila* samples from.** Three asterisks (***) denotes a highly significant result (p ≤ 0.001), based on paired Wilcoxon tests.

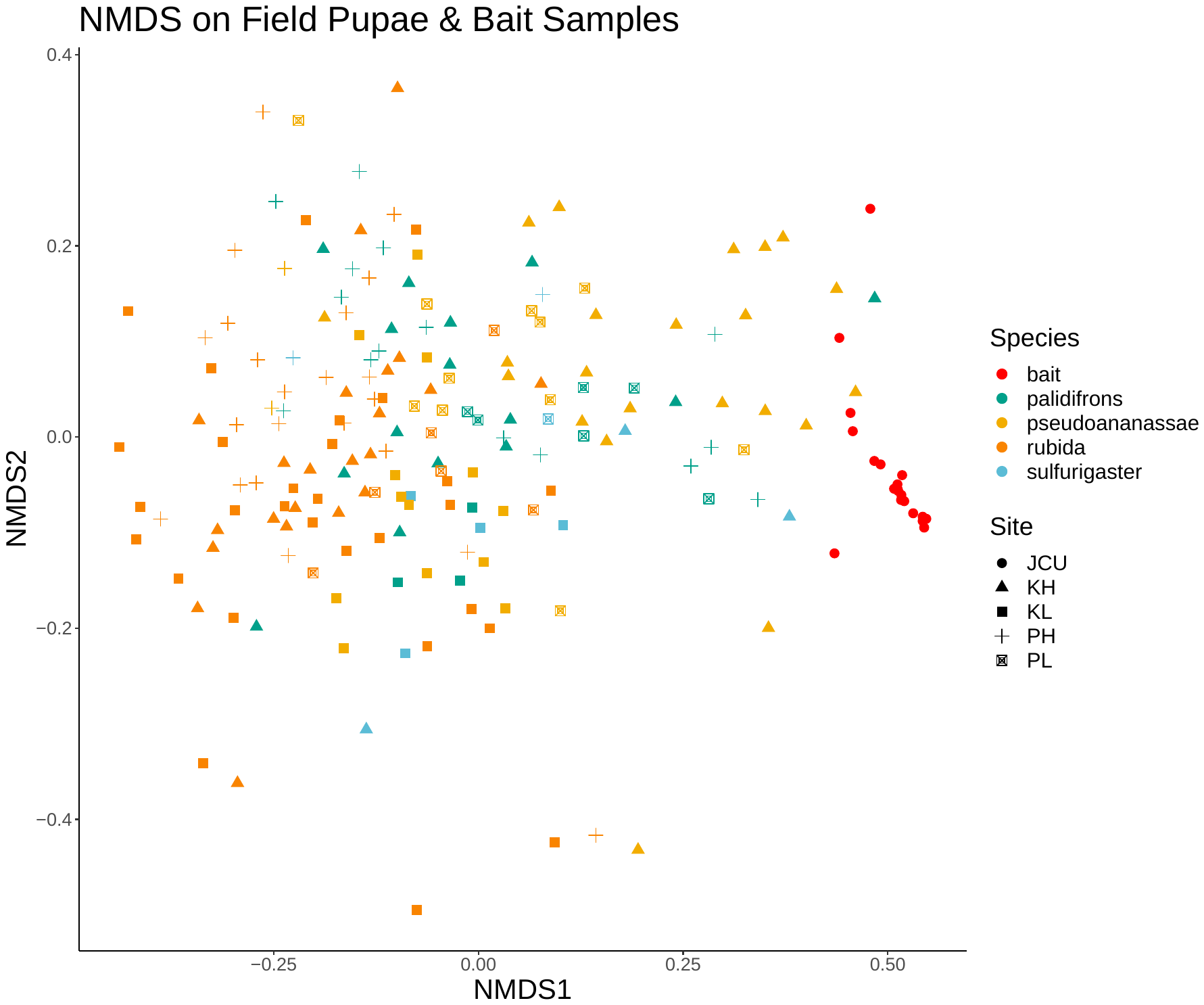

**Figure S13: NMDS of microbiome communities from pupae and bait samples originating from the field in this study. Symbols represent site of origin and different colours represent different species.**

| **Table S1: Summary statistics from PERMANOVA test on Bray-Curtis dissimilarity values for all samples from the lab and field combined. p values are corrected using the Benjamini-Hochberg method. Organised by F statistic in descending order.** | | | | | |
| --- | --- | --- | --- | --- | --- |
| **Response variable** | **Df** | **Sum of squares** | **R^2^** | **F statistic** | **p value** |
| Origin of sample | 1 | 20.68 | 0.15 | 107.52 | 0.001 |
| *Drosophila* species | 4 | 8.62 | 0.06 | 11.20 | 0.001 |
| Life stage | 1 | 2.02 | 0.01 | 10.52 | 0.001 |
| Field site | 5 | 9.44 | 0.07 | 9.82 | 0.001 |
| Gradient | 3 | 3.24 | 0.02 | 6.62 | 0.001 |
| Isofemale line | 12 | 8.84 | 0.06 | 3.83 | 0.001 |
| Trap exposure time | 5 | 4.71 | 0.03 | 3.50 | 0.001 |
| Trap identity | 15 | 10.25 | 0.07 | 3.33 | 0.001 |
| Generation number | 8 | 3.60 | 0.02 | 2.34 | 0.002 |

| **Table S2: Summary statistics from PERMANOVA test on Bray-Curtis dissimilarity values from core field samples. p values are corrected using the Benjamini-Hochberg method. Organised by F statistic in descending order.** | | | | | |
| --- | --- | --- | --- | --- | --- |
| **Response variable** | **Df** | **Sum of squares** | **R^2^** | **F statistic** | **p value** |
| Life stage | 1 | 4.56 | 0.07 | 17.61 | 0.001 |
| Field site | 2 | 2.03 | 0.07 | 9.82 | 0.001 |
| Trap exposure time | 1 | 2.11 | 0.03 | 8.74 | 0.001 |
| Elevation | 1 | 1.81 | 0.03 | 6.97 | 0.001 |
| Gradient:Site | 1 | 0.41 | 0.02 | 6.62 | 0.001 |
| *Drosophila* species | 3 | 3.68 | 0.05 | 4.73 | 0.001 |
| Gradient | 2 | 1.55 | 0.02 | 3.20 | 0.001 |
| Trap identity | 16 | 9.43 | 0.13 | 2.43 | 0.001 |
| Site:Trap | 17 | 8.87 | 0.13 | 2.16 | 0.021 |

| **Table S3: Summary output of DeSEQ analysis showing the top 5 most differentially abundant bacterial genera between host species sampled from the field.** | | | | | |
| --- | --- | --- | --- | --- | --- |
| Genus | baseMean | log2foldchange | lfcSE | Test statistic | Adjusted p value |
| ***Kozakia*** | 4.74 | -3.24 | 0.77 | -4.22 | ≤ 0.001 |
| ***Corynebacterium 1*** | 6.04 | -2.99 | 0.82 | -3.65 | ≤ 0.001 |
| ***Bacteroides*** | 23.33 | -5.90 | 1.03 | -5.74 | ≤ 0.001 |
| ***Marinospirillum*** | 25.31 | 3.53 | 0.97 | 3.33 | ≤ 0.001 |
| ***Morganella*** | 5.71 | -3.18 | 0.87 | -3.64 | ≤ 0.001 |

| **Table S4: Summary statistics from PERMANOVA test on Bray-Curtis dissimilarity values from lab samples. p values are corrected using the Benjamini-Hochberg method. Organised by F statistic in descending order.** | | | | | |
| --- | --- | --- | --- | --- | --- |
| **Response variable** | **Df** | **Sum of squares** | **R^2^** | **F statistic** | **p value** |
| Life stage | 1 | 1.91 | 0.06 | 15.79 | 0.001 |
| *Drosophila* species | 3 | 3.53 | 0.11 | 9.72 | 0.001 |
| IFLine:Generation | 4 | 2.95 | 0.09 | 6.09 | 0.001 |
| Isofemale Line | 6 | 3.95 | 0.13 | 5.44 | 0.001 |
| Generation | 8 | 3.66 | 0.12 | 3.79 | 0.001 |
| Field site | 2 | 0.89 | 0.03 | 3.71 | 0.001 |
| Elevation | 1 | 0.43 | 0.01 | 3.55 | 0.006 |
| Sex | 1 | 0.22 | 0.01 | 1.81 | 0.101 |
